# Predicting transfer RNA gene activity from sequence and genome context

**DOI:** 10.1101/661942

**Authors:** Bryan Thornlow, Joel Armstrong, Andrew Holmes, Russell Corbett-Detig, Todd Lowe

**Affiliations:** Department of Biomolecular Engineering, University of California, Santa Cruz, CA 95064; Genomics Institute, University of California, Santa Cruz, CA 95064

## Abstract

Transfer RNA (tRNA) genes are among the most highly transcribed genes in the genome due to their central role in protein synthesis. However, there is evidence for a broad range of gene expression across tRNA loci. This complexity, combined with difficulty in measuring transcript abundance and high sequence identity across transcripts, has severely limited our collective understanding of tRNA gene expression regulation and evolution. We establish sequence-based correlates to tRNA gene expression and develop a tRNA gene classification method that does not require, but benefits from comparative genomic information, and achieves accuracy comparable to molecular assays. We observe that guanine+cytosine (G+C) content and CpG density surrounding tRNA loci is exceptionally well correlated with tRNA gene activity, supporting a prominent regulatory role of the local genomic context in combination with internal sequence features. We use our tRNA gene activity predictions in conjunction with a comprehensive tRNA gene ortholog set spanning 29 placental mammals to infer the frequency of changes to tRNA gene expression among orthologs. Our method adds an important new dimension to tRNA annotation and will help focus the study of natural tRNA variants. Its simplicity and robustness enables facile application to other clades and timescales, as well as exploration of functional diversification of tRNAs and other large gene families.

## INTRODUCTION

Transfer RNAs (tRNAs) are essential for the translation of messenger RNA (mRNA) into proteins for all life. At the gene level in eukaryotes, they are of special interest for their high copy number, strong nucleotide sequence conservation, and variation in expression (Pan 2018; Kutter et al. 2011; Schmitt et al. 2014). tRNA molecules are required in large abundance to meet the dynamic metabolic needs of cells, and tRNA genes are believed to be among the most highly transcribed genes in the genome (Palazzo and Lee 2015; Boivin et al. 2018).

Despite high cellular demands, numerous individual tRNA genes have no direct evidence for expression (Kutter et al. 2011; Palazzo and Lee 2015; Hummel et al. 2019; Gogakos et al. 2017). High duplication rates and consequent weakened purifying selection may lead to an abundance of pseudogenes. Additionally, many of these genes may be tRNA-derived short interspersed nuclear elements (SINEs), which often retain strong promoter elements. However, even after removal of apparent pseudogenes and SINEs, more than 60 human tRNA genes and over 100 mouse tRNA genes have no epigenomic evidence for expression in any of more than 100 tissues and cell lines (Thornlow et al. 2018). Chromatin-immunoprecipitation sequencing (ChIP-Seq) data supports this conclusion, as one multi-species study detected expression of only 224 of 416 high-confidence tRNA genes in human liver, with other mammals showing similar patterns (Kutter et al. 2011).

tRNA gene expression may co-evolve with phenotypic differences between species. Data from prior studies suggests that the rate of evolution of protein coding gene expression levels differs by clade (Necsulea and Kaessmann 2014; Brawand et al. 2011; Li et al. 1996). The rate of evolution of gene expression also varies among non-coding RNA gene families (Meunier et al. 2013; Necsulea and Kaessmann 2014; Necsulea et al. 2014). Due to difficulties in high-throughput, accurate quantification of tRNA abundance, the complexity of tRNA gene expression across mammals is not well understood. The expanding functional repertoire of tRNA transcripts and fragments (Kirchner and Ignatova 2015; Goodarzi et al. 2015; Mleczko et al. 2014; Sun et al. 2018) indicates that changes in tRNA gene expression between species may have profound phenotypic effects.

Expression of tRNA genes has clear importance for organismal development and contribution to disease, but our understanding of its regulation and evolution is severely lacking for several reasons (Schaffer et al. 2014; Hanada et al. 2013; Yoo et al. 2016). Measuring expression of unique mRNA transcripts has become relatively straightforward. However, tRNA sequencing by methods originally developed for unmodified small RNAs (e.g., microRNAs) is frequently impeded by numerous RNA modifications at the reverse transcription phase. Only very recently have specialized sequencing library preparation methods been developed to remove or overcome these modifications, enabling effective sequencing (Cozen et al. 2015; Zheng et al. 2015). Furthermore, because the fully processed tRNA gene transcripts from different loci are often identical, simple tRNA-seq abundance measurements are not sufficient to determine the true transcriptional activity at each gene locus. Therefore, in order to determine which tRNA genes are potentially constitutively expressed, highly regulated, or silenced, other methods are needed.

Several genome-wide methods examine tRNA loci in their generally unique genomic contexts, bypassing the problem of identical mature tRNA transcripts. Such assays include chromatin immunoprecipitation (ChIP; Bogu et al. 2015; Roadmap Epigenomics Consortium et al. 2015; Thornlow et al. 2018), RNA Polymerase III (Pol III) ChIP-Seq (Kutter et al. 2011), and ATAC-Seq (Foissac et al. 2018), among others. These high-throughput assays are cost- and resource-intensive, so currently available data are often limited to few species and tissues. Importantly, these data show that identical tRNA genes do vary in expression profiles (Pan 2018; Kutter et al. 2011; Schmitt et al. 2014), supporting the need to incorporate extrinsic factors into the prediction of when or if tRNA genes are active. The study of the local genomic context is therefore essential, and has not been tackled comprehensively by any tRNA gene prediction method.

Here, we begin to resolve these concerns by developing a model to predict whether individual tRNA genes are actively transcribed, or largely silent. Prior work has shown that relative tRNA expression may be inferred based on DNA variation driven by transcription-associated mutagenesis (Thornlow et al. 2018). We leverage this correlation, further enhanced by other genomic features, to infer expression of tRNA genes with high accuracy. This novel advance in tRNA research effectively leverages, but does not require, comparative genomic information, enabling its broad application. We demonstrate our method using 29 placental mammalian genomes, most of which have no associated tRNA expression data. We also developed a robust mapping of syntenic tRNA gene orthologs across all 29 species. By combining our new method with this comprehensive ortholog set, we have developed, analyzed, and compared expression classifications over 10,000 tRNA genes, yielding a first look at the rate of tRNA gene regulation evolution in placental mammals, as well as the surprisingly high frequency of silenced “high-scoring” canonical tRNA genes.

## RESULTS

Our goal was to develop a tRNA “activity” predictive model that could be applied to as many species as possible. To date, the most facile method for inferring tRNA gene function has been the use of tRNAscan-SE covariance model bit scores, which quantify similarity to primary sequence and secondary structure profiles derived from an alignment of reference tRNAs. However, comparison of tRNAscan-SE bit scores to Pol III ChIP-Seq data from multiple mouse tissues suggests that high covariance model bit scores do not always correspond to expression (Kutter et al. 2011; Supplemental Fig. S1). To improve prediction of tRNA functional roles and better understand the basis of tRNA gene regulation in mammals, we evaluated many additional sequence features easily obtained from each gene’s sequence and genomic context. By quantifying the correlation of each of these features with gene activity from published epigenetic chromatin-based experimental data for human tRNA genes (Roadmap Epigenomics Consortium et al. 2015; Thornlow et al. 2018), we were then able to derive a model yielding accurate functional predictions on an independent test set (Fig. 1).

**Figure 1:**
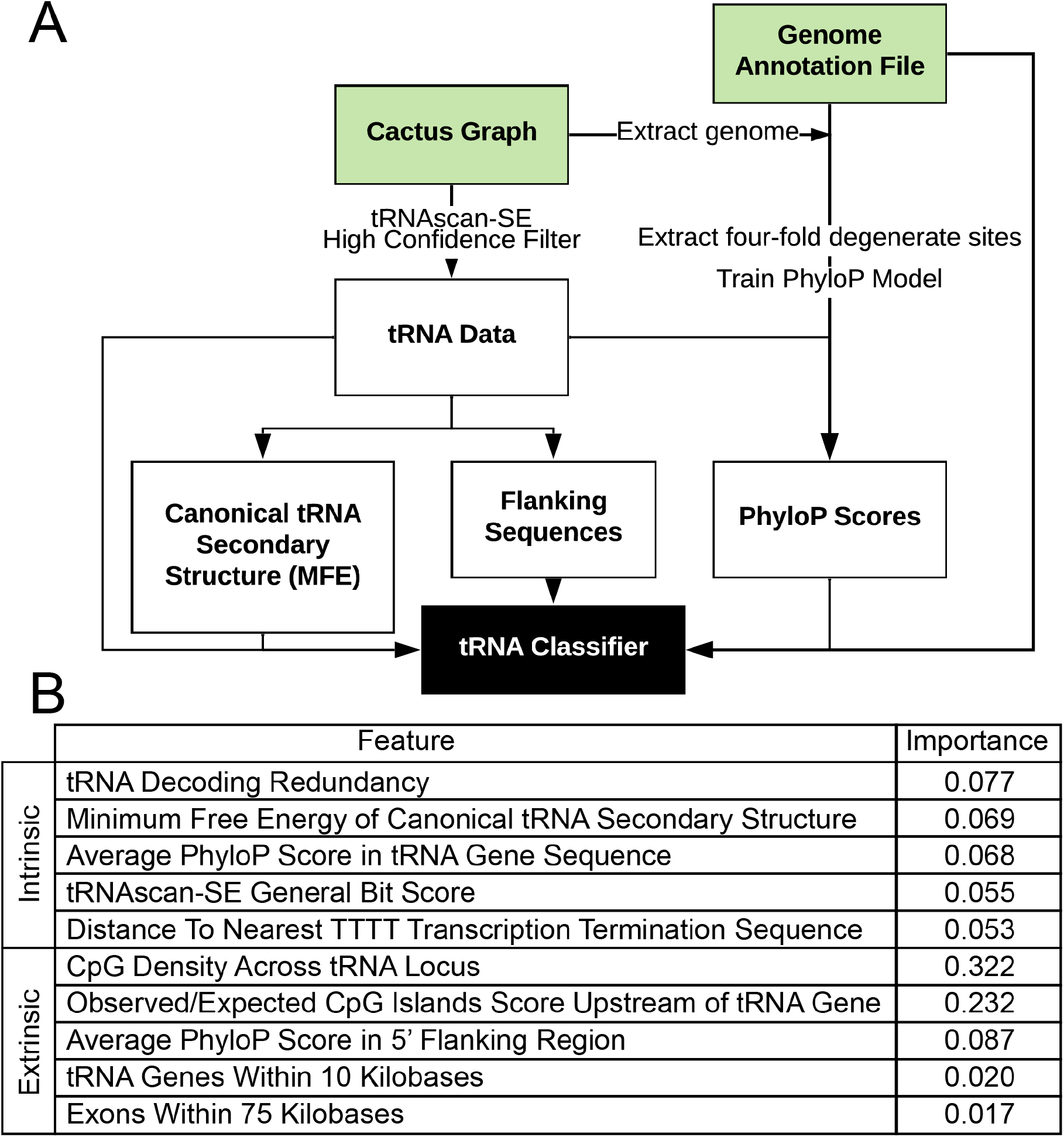
Both intrinsic (tRNA-specific) and extrinsic (genome context) features are integral to the model. (A) A schematic flowchart of our pipeline, and (B) all features included in the model with their relative importance values as measured by decrease in node impurity by scikit-learn (Pedregosa et al. 2011; see Methods). Green blocks indicate files not created by the pipeline. By default, our pipeline uses a gene annotation file and a Cactus graph (Armstrong et al. 2019; Paten et al. 2011a, 2011b; Nguyen et al. 2015), which is a reference-free whole genome alignment, as input. Higher importance scores indicate greater contribution to discrimination between active and inactive tRNA genes by the model. tRNA decoding redundancy refers to the number of tRNA genes sharing the same anticodon, and minimum free energy of canonical tRNA secondary structure refers to the minimum free energy when constrained to folding into the canonical cloverleaf structure (Lorenz et al. 2011). For calculating CpG-related statistics, we considered the tRNA locus to begin 350 bp upstream and end 350 bp downstream of each gene. To calculate the PhyloP score in the 5’ flanking region, we considered only the 20 bp immediately upstream of each tRNA gene.

To create our predictive model, we evaluated and incorporated two types of function-predictive statistics: intrinsic features related to tRNA gene sequence, and extrinsic features derived entirely from the genomic context. First, we reasoned that highly expressed tRNA genes should generally encode strong internal promoter sequences, and their transcripts must fold stably into the canonical tRNA structure. Both of these types of information are incorporated into tRNAscan-SE bit scores (Chan et al. 2019). Furthermore, our previous study found that tRNA gene conservation is highest for actively transcribed tRNA genes, presumably due to stronger purifying selection on required sequence features (Thornlow et al. 2018). Thus, we included tRNA gene conservation in the form of the PhyloP score, a nucleotide-level quantitative measure of conservation using multiple alignments (Pollard et al. 2010). We also assessed the correlation of gene activity with the length of each pre-tRNA’s 3’ tail, measured by the nucleotide distance from the end of the mature tRNA gene to the beginning of the “poly-T” transcription termination sequence (Allison and Hall 1985; Koski et al. 1980). Multiple studies on tRNA transcription termination (Maraia et al. 1994; Hamada et al. 2000; Orioli et al. 2011; Arimbasseri et al. 2013) observed that the RNase Z-trimmed 3’ sequences vary in overall length, composition, and terminator strength (poly-T length), each potentially affecting tRNA maturation and processing. We found that tRNAscan-SE bit scores, average PhyloP scores across tRNA gene sequences, and the distance to transcription termination sites are indeed significantly correlated with tRNA gene activity based on epigenomic data (Roadmap Epigenomics Consortium et al. 2015; Thornlow et al. 2018; Spearman rank correlation, p < 1e-5 for all comparisons).

Second, because mRNA expression depends heavily on local chromatin context, we explored features of the genomic environment. Protein coding genes in regions rich in CG dinucleotides (also known as CpGs) are known to be more frequently expressed (Gardiner-Garden and Frommer 1987; Krinner et al. 2014). Gardiner-Garden and Frommer define CpG island scores as the observed frequency of CpG dinucleotides compared to their expected frequency given the G+C content of a region. We found that these scores, when calculated for the 350 bases upstream of each gene, are significantly correlated with tRNA gene activity (Spearman rank correlation, p < 2.1e-26). Similarly, the frequency of CpG dinucleotides spanning from 350 bases upstream to 350 bases downstream of each tRNA gene is significantly correlated with tRNA gene activity (Roadmap Epigenomics Consortium et al. 2015; Thornlow et al. 2018; p < 8.0e-30). We also previously found that the putatively neutral regions immediately flanking highly expressed tRNA genes are more divergent, consistent with transcription-associated mutagenesis (Thornlow et al. 2018). Based on this, we found that the average PhyloP score of the 20-nucleotide 5’ flanking regions of tRNAs is significantly correlated with tRNA gene activity (Roadmap Epigenomics Consortium et al. 2015; Thornlow et al. 2018; p < 1.5e-16). Finally, based on an expectation for increased chromatin accessibility for tRNA genes near other active genes, we analyzed for gene neighbor proximity and found that tRNA genes are indeed more likely to be in an active chromatin state if near protein-coding genes (p < 4.7e-5) or other tRNA genes (p < 1.36e-6).

We hypothesized that some combination of both intrinsic and extrinsic features could enable accurate computational inference of potential for tRNA gene activity. To develop an integrated model, we tested several common frameworks, including random forest (RF), logistic regression, and support vector machines. The RF classifier was most effective, achieving the greatest area under the receiver operating characteristic curve (AUC, Fig. 2A-B) based on ten-fold cross-validation of human tRNA gene data and subsequent application, without retraining, to mouse tRNA gene data (see Methods). We developed our initial model with 30 features (Supplemental Table S1) and gradually reduced it to 10 (Fig. 1B), using the correlation between each feature and the activity labels from epigenomic data, as well as inter-correlations between features (Hall 1998; Hall et al. 2009; Roadmap Epigenomics Consortium et al. 2015; Thornlow et al. 2018).

**Figure 2:**
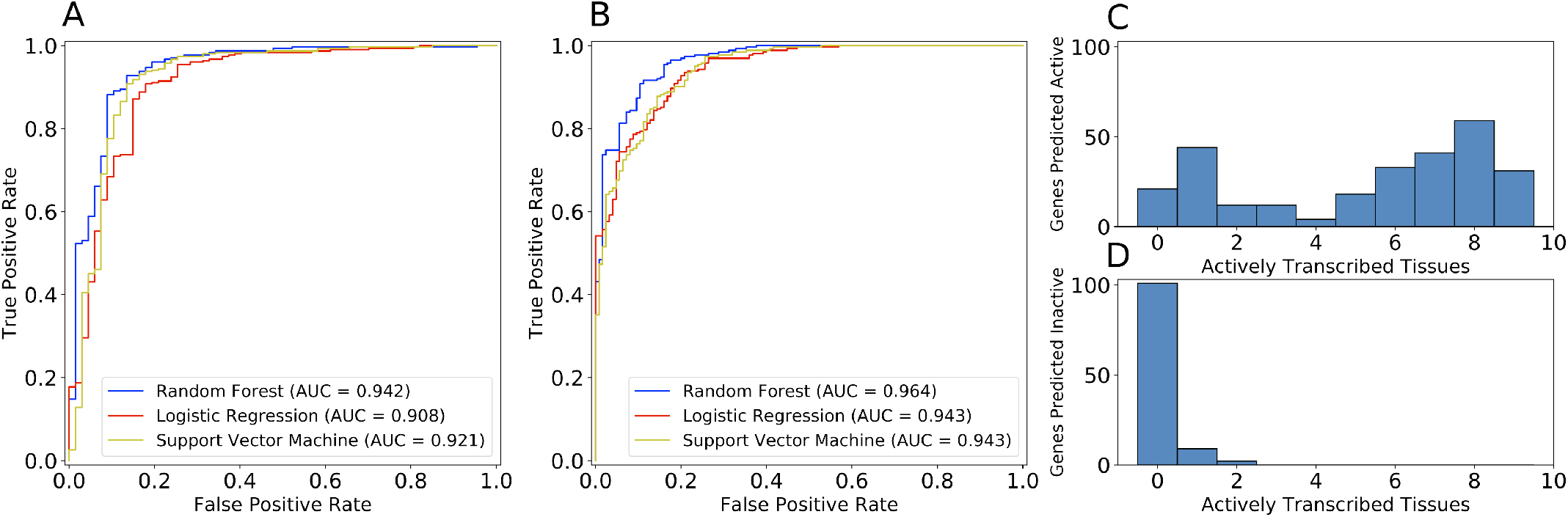
Random forest classifier achieves 92% accuracy on mouse tRNA genes. Receiver operating characteristic curves for random forest (blue), logistic regression (red) and support vector machine (yellow) upon application to (A) human training data with ten-fold cross-validation and (B) mouse test data. The number of mouse tRNA genes predicted (C) active and (D) inactive are compared to the number of tissues in which they are actively transcribed according to Bogu et al. 2015. We considered a mouse tRNA gene active if it is actively transcribed in at least one tissue.

### Features derived from CpG islands are most informative

To better understand and improve our classifier, we determined the relative importance of each feature, in terms of the normalized mean decrease in node impurity across all decision trees in the model (Fig. 1B; Pedregosa et al. 2011). We also used the Spearman rank correlation between each feature and the activity measurements to assess the strength of each feature. All features contribute to model accuracy and are significantly correlated with the activity labels (Spearman rank correlation, p < 1e-5 for all features), but the most informative features are derived from CpG islands at each tRNA locus, extolling the importance of chromosomal context and the ability to regulate expressed tRNA genes by altering methylation state (Gardiner-Garden and Frommer 1987).

One might expect that the tRNAscan-SE general bit score would be the most informative single feature (Fig. 1B, Table 1), but our analysis emphasizes the value of other measures, including the evolutionary conservation of the gene (PhyloP score), as well as that particular gene’s marginal importance among the full subset of tRNAs with the same anticodon (tRNA decoding redundancy). It is also important to note that we start with relatively high quality tRNAs with likely pseudogenes already removed by the tRNAscan-SE bit score-based high-confidence filter (Chan et al. 2019). Thus, for a starting set of tRNAs already vetted for reasonably strong features, the contribution of using the tRNAscan-SE bit score is marginally smaller than other features not previously used to estimate gene function. The high importance of CpG density in our results suggests that given a tRNA with a “passing” canonical set of features, based on tRNAscan-SE high-confidence predictions, it is the chromatin environment more than any other feature that determines tRNA gene activity potential (Fig. 1B, Table 1). Our classifier thus represents a substantial improvement over using tRNAscan-SE covariance bit scores alone (Supplemental Fig. S1).

**Table 1:** Our classifier uses all features to discriminate active and inactive tRNA genes. All human tRNA-Gln-TTG genes from our training data are shown with their feature values. Each column contains data for each gene locus, and the final two columns show the mean value across all active tRNA genes and all inactive tRNA genes in the training data. The header row follows GtRNAdb notation (Chan and Lowe 2016), with tRNA-Gln-TTG-3-1, 3-2 and 3-3 having identical mature gene sequences, distinct from tRNA-Gln-TTG-1-1, 2-1 and 4-1. All loci shown are correctly classified after ten-fold cross-validation.

|  | Gln-TTG-1-1 | Gln-TTG-2-1 | Gln-TTG-3-1 | Gln-TTG-3-2 | Gln-TTG-3-3 | Gln-TTG-4-1 | Active<br>Means | Inactive<br>Means |
| --- | --- | --- | --- | --- | --- | --- | --- | --- |
| **Activity** | **active** | **active** | **active** | **active** | **inactive** | **inactive** | - | - |
| tRNA Decoding<br>Redundancy | 6 | 6 | 6 | 6 | 6 | 6 | 11.1 | 18.4 |
| Minimum Free Energy<br>of Canonical tRNA<br>Secondary Structure | -27.1 | -27.1 | -22 | -22 | -22 | -24.1 | -27.4 | -23.7 |
| Average PhyloP Score<br>in tRNA Sequence | 1.635 | 0.457 | 0.517 | 0.641 | 1.051 | -0.047 | 0.856 | 0.335 |
| tRNAscan-SE General<br>Bit Score | 72.6 | 70 | 66.9 | 66.9 | 66.9 | 58.1 | 76.2 | 69.3 |
| Distance to Nearest<br>TTTT Transcription<br>Termination Sequence | 8 | 14 | 7 | 7 | 9 | 14 | 16.2 | 74.2 |
| CpG Density Across<br>tRNA Locus | 0.069 | 0.039 | 0.025 | 0.025 | 0.008 | 0.014 | 0.043 | 0.018 |
| Observed/Expected<br>CpG Islands Score<br>Upstream of tRNA<br>Gene | 0.753 | 0.492 | 0.624 | 0.571 | 0.147 | 0.111 | 0.67 | 0.27 |
| Average PhyloP Score<br>in 5' Flanking Region | 0.325 | -1.316 | -2.497 | -2.321 | -2.862 | 0.009 | -2.601 | -1.226 |
| tRNA Genes Within<br>10kb | 0 | 1 | 5 | 5 | 1 | 0 | 1.96 | 0.87 |
| Exons within 75kb | 40 | 25 | 37 | 37 | 15 | 0 | 35.2 | 21.4 |

To illustrate the utility of our model, we have provided the distributions of each feature for active versus inactive genes (Supplemental Fig. S2) and a table of component features for the human tRNA-Gln-TTG gene family (Table 1, Supplemental Fig. S3). Through the RF algorithm, our classifier creates 250 decision trees, each of which uses a different combination of features to split the gene set into genes predicted to be active and genes predicted to be inactive. Features with greater importance split the gene set into two groups that adhere more strongly to the activity labels obtained from epigenomic data (Fig. 1B; Pedregosa et al. 2011). It follows that genes predicted to be active should generally have feature values closer to the mean across all active tRNA genes than the mean across all inactive tRNA genes from the training data. For example, tRNA-Gln-TTG-1-1, which is predicted to be active, is very well conserved (1.635 PhyloP score; Table 1), has a high tRNAscan-SE general bit score (72.6), is in a very CpG-rich region (0.753 observed/expected CpG islands score), and is near to many exons (40 within 75 kb). Conversely, tRNA-Gln-TTG-4-1, which is predicted to be inactive, is very weakly conserved as indicated by its low PhyloP score (−0.047), and has a low tRNAscan-SE general bit score (58.1; Chan et al. 2019) consistent with inactivity. Similarly, while tRNA-Gln-TTG-3-3 has an exactly identical nucleotide sequence as tRNA-Gln-TTG-3-1 and 3-2, tRNA-Gln-TTG-3-3 is proximal to fewer protein coding exons (15 within 75 kb) and tRNA genes (1 within 10 kb), has a more distant poly-T transcription termination sequence (9 nucleotides from the end of the gene), and is located in a less CpG-dense region (0.008 CpG dinucleotides / total dinucleotides). Based on the characteristics of these features as described in these examples, our classifier can accurately discern tRNA gene activity.

### Our classifier is 92% accurate in classifying mouse tRNA genes

We tested the accuracy of our classifier using epigenomic and Pol III ChIP-Seq data for mouse genes. Our mouse tRNA gene set contains 387 genes, with 262 observed as active and 125 believed silent based on published experimental data (Bogu et al. 2015; see Methods). Our classifier predicted that 275 of these genes are active and that 112 are inactive, correctly categorizing 356 tRNA genes and achieving 92.0% accuracy (Fig. 1C-D). Of the 31 misclassified mouse tRNA genes, 22 are misclassified as active and 9 are misclassified as inactive. We note that these genes are not biased by isotype (Supplemental Table S2), nor by genomic location, and are therefore most likely misclassified for a variety of reasons. Of the 22 misclassified as active, some of these may reflect incompleteness of cell types or developmental states in the reference epigenomic data. Some misclassified tRNA genes may also serve important specialized functions not fully captured by our feature set, such as acting as chromatin insulator elements (Raab et al. 2012).

### Classification without alignment or annotation is similarly accurate

We developed our method such that it could potentially be applied to any species with a sequenced genome. For best performance, we used a Cactus graph (Armstrong et al. 2019; Paten et al. 2011a, 2011b; Nguyen et al. 2015), which is a reference-free whole genome alignment, incorporating 29 genomes (Supplemental Table S3), but we recognize that Cactus graphs are not yet available for all species. To accommodate species for which no alignments or protein-coding gene annotations have been developed, we included an option to remove features relying on these inputs. Use of this model led to decreases in accuracy in both human (AUC = .932, 92.5% accuracy compared to AUC = .942 and 93.0% accuracy in the full model) and mouse (AUC = .935, 90.1% accuracy compared to AUC = .964 and 92.0% accuracy in the full model), which may be exacerbated upon application to more phylogenetically distant species.

### ChIP-Seq and ATAC-Seq data further validate our classifications in additional species

To further test our model, we compared our predictions to RNA Polymerase III (Pol III) ChIP-Seq data previously collected from the livers of four species (*Mus musculus, Macaca mulatta, Rattus norvegicus* and *Canis lupus familiaris*; Kutter et al. 2011). We found roughly expected agreement between our classifications and the Pol III ChIP-Seq read counts from a single tissue (Fig. 3). We also found that the relationship between the probability scores in our classifications and the Pol III ChIP-Seq read counts is consistent across all four species (i.e., a similar majority of predictions are far left or far right in Fig. 3), suggesting similar accuracy in classification. Our predictions are also similarly accurate when compared to mouse muscle and testes ChIP-Seq data (Supplemental Fig. S4).

**Figure 3:**
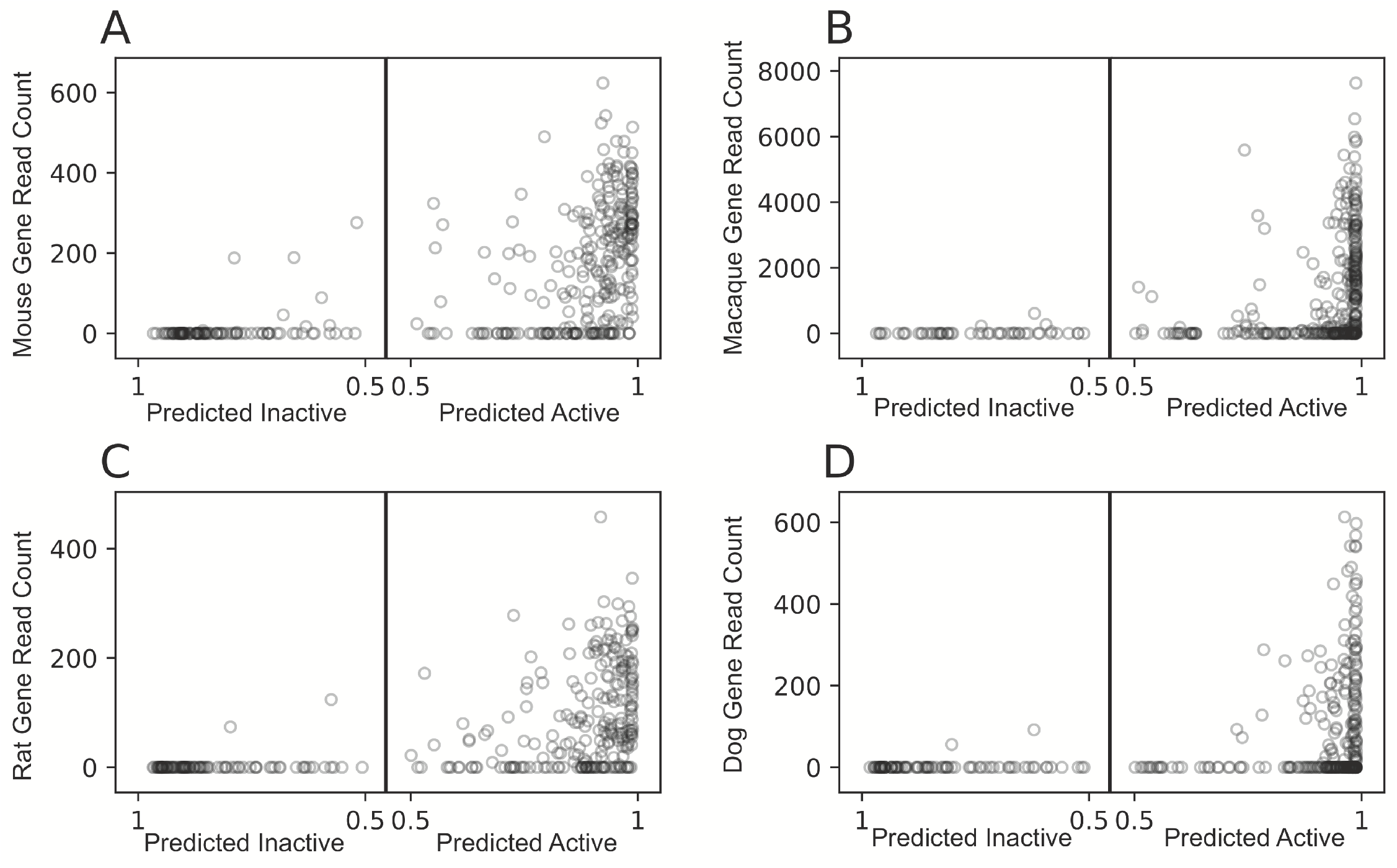
Classification of gene activity based on genomic data achieves similar results to ChIP-Seq expression analysis in four species. Probability scores output by the classifier for (A) mouse, (B) macaque, (C) rat and (D) dog are shown on the x-axis where tRNA genes further left are predicted inactive with greater probability, and tRNA genes further right are predicted active with greater probability. The y-axis shows RNA Polymerase III ChIP-Seq read counts from the liver of each species for each tRNA gene, taken from Kutter et al. 2011.

We predicted many tRNA genes as active despite a lack of Pol III binding at these loci in liver, muscle, and testes. This is not surprising, as our model does not predict expression in specific tissues, but is instead trained to predict tRNA genes as active if epigenomic data indicates active transcription in at least one of many tissues (Roadmap Epigenomics Consortium et al. 2015). For example, in mouse, 262 total tRNA genes are active in at least one tissue based on the epigenomic data, but 89 of these (34%) are not expected to be active in the liver based on the same data (Bogu et al. 2015). Our model predicted 122 (macaque), 68 (rat) and 149 (dog) tRNA genes as active despite Pol III ChIP-Seq read counts of zero in the liver (Kutter et al. 2011). Although ChIP-Seq has not been performed on macaque, rat and dog tRNA loci for other tissues, we found that virtually all tRNA genes with measured Pol III binding were predicted to be active by our classifier. Among tRNA genes with Pol III ChIP-Seq read-counts greater than zero, we predicted that 95.3% are active in mouse, 97.7% in macaque, 98.9% in rat and 98.5% in dog. This consistency in tRNA distributions and classifier behavior across species suggests that the classifier is similarly accurate in mouse, macaque, rat and dog.

To further validate our predictions, we used ATAC-seq data (Foissac et al. 2018), captured in liver, CD4 and CD8 cells for the cow, pig and goat genomes. We compared our predictions to the ATAC-Seq peaks across these tissues for the regions spanning from 250 base pairs upstream to 250 base pairs downstream of each tRNA gene (Supplemental Fig. S5). Due to the inclusion of only a small subset of tissues in this data, many tRNA loci that do not show activity in these tissues but were predicted active by our model may be active in other tissues. Among tRNA genes with ATAC-Seq peaks greater than zero, we predicted 90.4%, 95.1%, and 90.8% as active in cow, goat and pig, respectively. These results are comparable to measurements obtained from ChIP-Seq data in mouse, macaque, rat and dog. Decreases in accuracy may be due to differences in resolution between ATAC-Seq and ChIP-Seq, or increased phylogenetic distance from human to these species, relative to mouse, macaque, rat and dog.

### tRNA gene classifications follow similar distributions across the eutherian phylogeny

We applied our model to 29 mammalian species to glean new insights into the evolution of tRNA genes (Fig. 4, Supplemental Table S3). We determined the distributions of active and inactive tRNA genes by anticodon across these species, finding that most species have approximately 250-350 predicted active genes, comprising roughly 75% of their tRNA gene sets (Fig. 4, Supplemental Table S4). We observed similar distributions by clade, with few exceptions. Notably, *Bos taurus, Capra hircus* and *Orcinus orca* (cow, goat and orca, respectively) have more than 300 predicted inactive tRNA genes while no other species has more than 156. This may reflect decreased ability of tRNAscan-SE to discriminate tRNA-derived SINEs (short interspersed nuclear elements) from tRNA genes in these species (Chan et al. 2019). Consistent with Tang et al. 2009, we identified 406 active cow tRNA genes, or 437 when including those found in segmental duplications (Supplemental Table S5; see Methods), the most of any species in our study. Of the 439 “core” cow tRNA genes identified by Tang et al. 2009, there are 396 genes with exactly matching sequences in our high confidence tRNA gene set (Chan et al. 2019), and we classified 346 of these as active. Gene sequences found by Tang et al. 2009 that were not in our set are likely due to differences in updated genome assemblies and improvements in tRNAscan-SE 2.0 (Chan et al. 2019).

**Figure 4:**
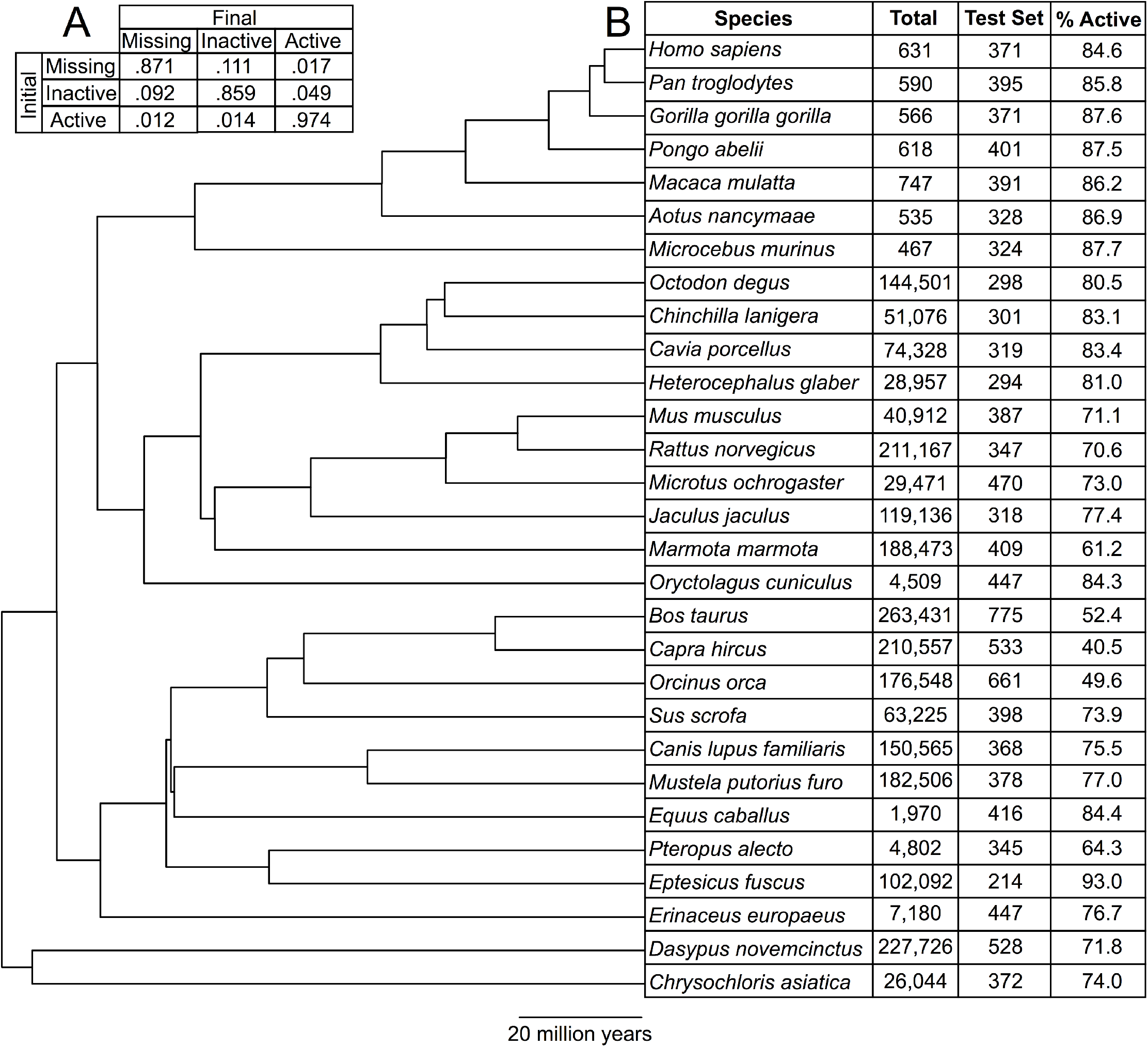
Changes between activity states are rare among placental mammal tRNA genes. (A) We estimated transition probabilities between each predicted activity state over a branch length of 10 million years using RevBayes (Höhna et al. 2016). “Missing” refers to cases where no syntenic ortholog is present for a given species. (B) For each species in our phylogeny (Hedges et al. 2006), the number of tRNA genes annotated by tRNAscan-SE 2.0 (Total; Chan et al. 2019), the number of tRNA genes present after application of the high-confidence filter and removal of tRNA genes in segmental duplications (Test Set; see Methods), and the percent of tRNA genes in the test set classified as active (% Active), are shown. For human and mouse, tRNA genes with no epigenomic data are excluded from this table as well (see Methods).

We verified that all species had at least one tRNA gene predicted as active for each expected anticodon (Grosjean et al. 2010, Supplemental Table S4). This excludes anticodons with no known active tRNA genes in human or mouse. We observed only three exceptions to this rule, which most likely represent genome assembly or classification errors (see Supplemental Material).

### Deeply conserved tRNA genes are more likely to be active

In order to investigate the relationship between evolutionary conservation and transcriptional activity, we developed a complete set of placental mammal tRNA gene orthologs using a Cactus graph (Armstrong et al. 2019; Supplemental Tables S3, S6). Cactus graphs are state-of-the-art alignments that allow greater detection of synteny across many species. Of the 11,752 tRNA genes in our 29-species alignment, 3,582 genes in total, or about 122 per species on average, appear to be species-specific. We condensed the remaining 8,170 genes into 1,097 ortholog sets. On average, each of these orthologs sets spans 7.4 species, indicating that tRNA genes are generally either fairly deeply conserved or recently evolved (Supplemental Fig. S6). This is consistent with prior studies in *Drosophila* showing that tRNA genes can be “core” or “peripheral” (Rogers et al. 2010).

We identified a “core” set of 97 primate tRNA genes (Supplemental Fig. S7), for which all seven primate species (human, chimpanzee, gorilla, orangutan, macaque, *Microcebus murinus* (gray mouse lemur) and *Aotus nancymaae* (Nancy Ma’s night monkey)) have a syntenic ortholog. These most likely represent tRNA genes present in the primate common ancestor that have not been lost in any lineage leading to the sampled genomes. These genes encode 19 amino acids (Supplemental Fig. S8). Surprisingly, a single standard amino acid isotype is not represented: cysteine. tRNA-Cys genes are often present in high numbers, and every species in the primate phylogeny has at least 17 such genes. However, tRNA-Cys genes are prone to accumulating nucleotide substitutions, and the human genome contains 23 unique high-confidence tRNA-Cys-GCA gene sequences, the most of any isotype. Therefore, the lack of a “core” eutherian tRNA-Cys gene may be due to relatively rapid evolution of this gene family, or perhaps difficulty in alignment due to their high variation in sequence.

In 15 of these 97 “core” ortholog sets, we predicted at least one member of the ortholog set as active and at least one as inactive among the different primate species. Across all 97 “core” ortholog sets, we predict 98% of all member tRNA genes as active, suggesting that deeply conserved tRNA genes are highly likely to be active. Out of all 1,097 ortholog sets in placental mammals, 750 contain only tRNA genes predicted to be active, approximately mirroring the distribution of active to inactive tRNA genes predicted at the species level (Fig. 4B).

### Substitutions in tRNA anticodons accompany changes in predicted activity state

Among our 1,097 ortholog sets, we identified 113 sets containing genes corresponding to multiple different anticodons. 53 of these 113 anticodon shifts (46%) are non-synonymous, resulting in a change in the corresponding amino acid. Ortholog sets containing anticodon shifts are significantly enriched for changes in predicted activity, as 37 of the 113 ortholog sets contain at least one predicted active and at least one predicted inactive tRNA gene (Hypergeometric test, p < 1.5e-4). Likewise, of the 53 ortholog sets containing non-synonymous anticodon substitutions, 19 contain at least one predicted active and at least one predicted inactive tRNA gene (Hypergeometric test, p < 1.4e-3).

### Transitions between active and inactive are rare

We fit our ortholog sets and their predicted activity states to a Markov model of evolution of discrete characters using RevBayes (Höhna et al. 2016; Fig. 4A; see Methods). By fitting our data to the model, we estimated transition probabilities to and from three states - active, inactive, and missing (no detected ortholog). We held the phylogeny constant and solved only for the transition rate parameters. Consistent with the premise that local gene duplication is the dominant mode of tRNA gene evolution, our model suggests that active tRNA genes overwhelmingly tend to remain active, and inactive tRNA genes tend to remain inactive (Fig. 4A).

Although activity state transition events are rare, our classifier does well to detect them. There are 188 human/mouse ortholog pairs for which we have epigenomic data, and in 173 (92%) of them, human and mouse have the same activity state based on the epigenomic data. However, we correctly classified 183 human (97%) and 181 mouse (96%) tRNA genes within this set, indicating that our classifier detected activity state changes between these species. Assuming that the activity state of orthologous tRNA genes remains constant across closely related species would yield largely accurate activity state predictions for annotating tRNAs in additional new species. However, our classifier represents an improvement over this assumption, and is also applicable to species-specific tRNA genes, which are especially common and have no ortholog data.

Overall, transitions in activity state of any kind are rare, with the active state being most stable among orthologs (Fig. 4A). Surprisingly, inactive tRNA genes that are conserved most often remain inactive as well (Fig. 4A), hinting at important biological roles for conserved, apparently silent tRNA genes. We also observed some variation in the relative transition probabilities within clades (Supplemental Fig. S9). Primate tRNA genes are less likely to remain in their initial predicted activity state than rodent tRNA genes. This is consistent with prior studies on the rate of evolutionary change of protein coding gene expression between clades (Brawand et al. 2011; Necsulea and Kaessmann 2014) but most likely reflects differences in sample size between clades. Based on our results, turnover in tRNA gene expression class generally appears to be slow, similar to protein coding genes (Brawand et al. 2011).

## DISCUSSION

Greater understanding of tRNA regulation is a difficult and unmet challenge. There are many obstacles preventing direct measurement of expression at the gene level. Nonetheless, we have shown that, in order to determine the transcriptional potential of tRNA genes, direct measurement across many tissues is not necessarily required if the gene sequence and genomic context is known. We leverage features both intrinsic to tRNA genes, which relate directly to tRNA function and processing, and extrinsic, which relate to the chromosomal region regulation.

Our classifier demonstrates remarkable accuracy in predicting tRNA gene activity. However, there are caveats to our method. Importantly, the epigenomic chromatin-IP data used to create the “active” and “inactive” gene training data for both human and mouse data survey many, but by no means all cell types. It is possible that many tRNA genes that are inactive according to the epigenomic data are in fact very tightly regulated, such that their activity is not captured by prior studies. For example, a subset of apparently silent human tRNA genes are present in the introns of protein coding genes, and may regulate their transcription by low-level transcriptional interference (Yeganeh et al. 2017). Additionally, while chromatin-IP captures the chromosomal environment across many tissues and is by far the most comprehensive data available for estimating individual tRNA gene activity, it is only available for a few species. The rarity of comprehensive tRNA gene expression data motivates the creation of our classifier. Available data indicate that our estimates are comparable to experimental results, but with much greater ease of use and cost-effectiveness.

Our approach has broad applicability. We have shown that accounting for the genomic context allows for much improved annotation of tRNA genes, as high copy-number and high sequence identity have previously made it difficult to determine which tRNA genes are indeed actively expressed. We have focused on tRNA genes, but the basic principles underlying our model might be applied to any multi-copy gene family. For example, we have previously shown that histone protein coding genes exhibit similar flanking genetic variation to tRNA genes (Thornlow et al. 2018). A model similar to ours could be developed to infer relative levels of expression of histone protein coding genes.

Our methods may also be applied to other species. We have previously demonstrated that transcription-associated mutagenesis creates signatures at plant and insect tRNA gene loci similar to those in human and mouse (Thornlow et al. 2018). While some features may not extend quite so far phylogenetically, and others may require recalibration or processing, a similar model could be developed for other eukaryotic clades. We chose to work with placental mammals for this study, as the corresponding data is of relatively high quality.

Classification using our approach may expand in several directions as more data is collected. For example, variation within populations may be useful for predicting relative transcript expression within gene families. We previously determined that actively transcribed tRNA genes accumulate more rare single nucleotide polymorphisms (SNPs) in both their flanking regions and gene sequences (Thornlow et al. 2018). Therefore, we expect that when population variation data is available for more species, we may infer expression differences at narrower timescales. Our model may also be expanded to accommodate non-binary classification of expression levels in different tissue types, and better capture the nuance of tRNA gene expression regulation.

In the future, annotations created by our method will be useful in prioritizing tRNA characterization experiments, as well as interpreting the biological effects of mutations in and surrounding tRNA genes. This work informs the broader questions of regulation and evolution of tRNA genes in mammals, quantitatively illustrating that tRNA gene expression is dependent on features intrinsic to the gene sequence as well as their rich diversity of genomic environments.

## METHODS

### Developing and testing the classifier

For the training data, we used coordinates from human genome assembly GRCh38 for tRNA genes not removed by the tRNAscan-SE high confidence filter (Chan and Lowe 2016; Chan et al. 2019). For all species, including human and mouse, we extracted the genomes from our Cactus alignment (Armstrong et al. 2019; Supplemental Table S3), ran tRNAscan-SE 2.0, and applied the EukHighConfidenceFilter to exclude tRNA pseudogenes and tRNA-derived SINEs (Chan et al. 2019). We used custom Python scripts to find tRNA loci that were identical from 80 nucleotides upstream to 40 nucleotides downstream of the gene start and end. We considered these to be segmental duplications and excluded them from training and testing of the classifier. If any of these loci also did not align to any tRNA loci in any other species, they were also removed from our ortholog sets, as they most likely represent assembly errors. For genomes in which at least 85% of nucleotides were found on chromosomes, we excluded all tRNA genes not found on chromosomes. For the human tRNA gene set, because our epigenomic data is based on GRCh37 assembly gene annotations, we removed any tRNA genes that were not included in the older assembly (determined by performing liftOver (Casper et al. 2018) conversion from GRCh38 to GRCh37), as well as genes in segmental duplications in either assembly.

We used the PHAST (Hubisz et al. 2011) and HAL (Hickey et al. 2013) toolkits to generate PhyloP data, and RNAfold (Lorenz et al. 2011) to estimate minimum free energy, using the constraints on secondary structure output by tRNAscan-SE. We used custom Python scripts in conjunction with tRNAscan-SE output files and genome annotation files (accession numbers listed in Supplemental Table S3) to obtain data for all other features. When PhyloP data were unobtainable due to lack of alignment, we replaced feature values for each tRNA gene with the mean value for that feature across all tRNA genes in that species, using the SimpleImputer() module in scikit-learn (Pedregosa et al. 2011). We used scikit-learn to train the model and classify each gene (Pedregosa et al. 2011). We used the Spearman rank correlation test to ensure that no features were perfectly correlated (Guyon and Elisseeff 2003). We used CfsSubsetEval (Hall et al. 2009) to remove uninformative features and scikit-learn to determine feature importance (Pedregosa et al. 2011). See Supplemental Methods for more details.

To train and test our model, we used epigenomic data from the NIH Roadmap Epigenomics Program (Roadmap Epigenomics Consortium et al. 2015) and the chromatin state-associated gene study in mice (Bogu et al. 2015) for human and mouse tRNA gene activity states, respectively. These studies used histone marks to identify regions of active transcription across 127 human tissues and 9 mouse tissues, respectively. In both species, we excluded tRNA genes for which epigenomic data was not available, and tRNA genes contained within large segmental duplications as defined above. Our training set includes 371 human tRNA genes, 304 active and 67 inactive. For both species, we considered tRNA loci as active if they had an open chromatin state in at least one tissue. We considered all others inactive. To validate our model, we compared our classifications to ChIP-Seq read counts taken directly from Kutter et al 2011 and ATAC-Seq peaks taken directly from Foissac et al 2018, using liftOver (Casper et al. 2018) conversion to accommodate differences in genome assembly.

### Creating an ortholog set

We used hal2maf to create 29-way alignments for all tRNA loci of interest for the species in our phylogeny (Hickey et al. 2013). For each tRNA locus, we considered the best aligning tRNA locus from all other species as orthologous, allowing only one ortholog per species per locus. We augmented our ortholog sets with syntenic human/mouse, human/dog and human/macaque tRNA gene ortholog pairs from Holmes 2018. For all instances in which each tRNA gene in a Holmes 2018 ortholog pair aligned to mutually exclusive sets of species in our Cactus graph, we combined them into one ortholog set. We found that 29 Holmes 2018 human-mouse ortholog pairs align to each other in the Cactus graph, 152 align to mutually exclusive sets of species, and 17 align to overlapping sets of species. Therefore, we combined the 152 human-mouse tRNA gene pairs into larger ortholog sets.

### Fitting a Markov model

We entered our species into TimeTree (Kumar et al. 2017) to generate a phylogeny, and fit our phylogeny, orthologs sets and predicted activity states to a Markov model using RevBayes (Höhna et al. 2016). We held the phylogeny constant and allowed RevBayes to optimize only the Q matrix using our tRNA data. We then determined transition probabilities over 10 million years using the RevBayes function getTransitionProbabilities() across all species and by clade (Supplemental Fig. S9). See Supplemental Methods for more details.

## DATA ACCESS

Our pipeline is available at https://github.com/bpt26/tRNA_classifier/. Classification and ortholog data are available in Supplemental Tables S4-S6. Alignments generated by the Cactus graph are available upon request. Genome assemblies included in the Cactus graph are listed in Supplemental Table S3.

## ACKNOWLEDGMENTS

We thank the R.B.C.-D. and T.M.L. laboratories for suggestions and feedback. This work was supported by NIH/National Human Genome Research Institute Grant Award R01HG006753 (to T.M.L.) and NIH/National Institute of General Medical Sciences Grant Award R35GM128932 (to R.B.C.-D.). B.P.T. was funded by NIH/National Human Genome Research Institute Grants T32HG008345 and F31HG010584.

## Author contributions

Study design was by B.T., R.B.C.-D. and T.M.L.. J.A. created and provided the Cactus graph. A.H. provided mouse tRNA gene activity labels and ortholog data. B.T. wrote pipeline for extracting feature data and classifying genes. B.T., R.B.C.-D. and T.M.L. wrote the manuscript.

## DISCLOSURE DECLARATION

We declare no competing interests.

## Notes

#### Summary of Updates

Updates to wording and structure throughout manuscript and changes made to all main figures

## REFERENCES

Allison DS, Hall BD. 1985. Effects of alterations in the 3’ flanking sequence on in vivo and in vitro expression of the yeast SUP4-o tRNATyr gene. EMBO J 4: 2657–2664.

Arimbasseri AG, Rijal K, Maraia RJ. 2013. Transcription termination by the eukaryotic RNA polymerase III. Biochim Biophys Acta 1829: 318–330.

Armstrong J, Hickey G, Diekhans M, Deran A, Fang Q, Xie D, Feng S, Stiller J, Genereux D, Johnson J, et al. 2019. Progressive alignment with Cactus: a multiple-genome aligner for the thousand-genome era. http://dx.doi.org/10.1101/730531.

Beier H, Grimm M. 2001. Misreading of termination codons in eukaryotes by natural nonsense suppressor tRNAs. Nucleic Acids Res 29: 4767–4782.

Blanchette M, Kent WJ, Riemer C, Elnitski L, Smit AFA, Roskin KM, Baertsch R, Rosenbloom K, Clawson H, Green ED, et al. 2004. Aligning multiple genomic sequences with the threaded blockset aligner. Genome Res 14: 708–715.

Bogu GK, Vizán P, Stanton LW, Beato M, Di Croce L, Marti-Renom MA. 2015. Chromatin and RNA Maps Reveal Regulatory Long Noncoding RNAs in Mouse. Mol Cell Biol 36: 809–819.

Boivin V, Deschamps-Francoeur G, Couture S, Nottingham RM, Bouchard-Bourelle P, Lambowitz AM, Scott MS, Abou-Elela S. 2018. Simultaneous sequencing of coding and noncoding RNA reveals a human transcriptome dominated by a small number of highly expressed noncoding genes. RNA 24: 950–965.

Brawand D, Soumillon M, Necsulea A, Julien P, Csárdi G, Harrigan P, Weier M, Liechti A, Aximu-Petri A, Kircher M, et al. 2011. The evolution of gene expression levels in mammalian organs. Nature 478: 343–348.

Casper J, Zweig AS, Villarreal C, Tyner C, Speir ML, Rosenbloom KR, Raney BJ, Lee CM, Lee BT, Karolchik D, et al. 2018. The UCSC Genome Browser database: 2018 update. Nucleic Acids Res 46: D762–D769.

Chan PP, Lin BY, Mak AJ, Lowe TM. 2019. tRNAscan-SE 2.0: Improved Detection and Functional Classification of Transfer RNA Genes. http://dx.doi.org/10.1101/614032.

Chan PP, Lowe TM. 2016. GtRNAdb 2.0: an expanded database of transfer RNA genes identified in complete and draft genomes. Nucleic Acids Res 44: D184–9.

Cozen AE, Quartley E, Holmes AD, Hrabeta-Robinson E, Phizicky EM, Lowe TM. 2015. ARM-seq: AlkB-facilitated RNA methylation sequencing reveals a complex landscape of modified tRNA fragments. Nat Methods 12: 879–884.

Foissac S, Djebali S, Munyard K, Villa-Vialaneix N, Rau A, Muret K, Esquerre D, Zytnicki M, Derrien T, Bardou P, et al. 2018. Livestock genome annotation: transcriptome and chromatin structure profiling in cattle, goat, chicken and pig. bioRxiv 316091. https://www.biorxiv.org/content/early/2018/05/11/316091.abstract (Accessed November 20, 2018).

Gardiner-Garden M, Frommer M. 1987. CpG islands in vertebrate genomes. J Mol Biol 196: 261–282.

Gogakos T, Brown M, Garzia A, Meyer C, Hafner M, Tuschl T. 2017. Characterizing Expression and Processing of Precursor and Mature Human tRNAs by Hydro-tRNAseq and PAR-CLIP. Cell Rep 20: 1463–1475.

Goodarzi H, Liu X, Nguyen HCB, Zhang S, Fish L, Tavazoie SF. 2015. Endogenous tRNA-Derived Fragments Suppress Breast Cancer Progression via YBX1 Displacement. Cell 161: 790–802.

Grosjean H, de Crécy-Lagard V, Marck C. 2010. Deciphering synonymous codons in the three domains of life: co-evolution with specific tRNA modification enzymes. FEBS Lett 584: 252–264.

Guyon I, Elisseeff A. 2003. An Introduction to Variable and Feature Selection. J Mach Learn Res 3: 1157–1182.

Hall MA. 1998. Correlation-based feature subset selection for machine learning. Thesis submitted in partial fulfillment of the requirements of the degree of Doctor of Philosophy at the University of Waikato. https://ci.nii.ac.jp/naid/10018668219/.

Hall M, Frank E, Holmes G, Pfahringer B, Reutemann P, Witten IH. 2009. The WEKA data mining software: an update. ACM SIGKDD Explorations Newsletter 11: 10–18.

Hamada M, Sakulich AL, Koduru SB, Maraia RJ. 2000. Transcription termination by RNA polymerase III in fission yeast. A genetic and biochemically tractable model system. J Biol Chem 275: 29076–29081.

Hanada T, Weitzer S, Mair B, Bernreuther C, Wainger BJ, Ichida J, Hanada R, Orthofer M, Cronin SJ, Komnenovic V, et al. 2013. CLP1 links tRNA metabolism to progressive motor-neuron loss. Nature 495: 474–480.

Hedges SB, Dudley J, Kumar S. 2006. TimeTree: a public knowledge-base of divergence times among organisms. Bioinformatics 22: 2971–2972.

Hickey G, Paten B, Earl D, Zerbino D, Haussler D. 2013. HAL: a hierarchical format for storing and analyzing multiple genome alignments. Bioinformatics 29: 1341–1342.

Höhna S, Landis MJ, Heath TA, Boussau B, Lartillot N, Moore BR, Huelsenbeck JP, Ronquist F. 2016. RevBayes: Bayesian Phylogenetic Inference Using Graphical Models and an Interactive Model-Specification Language. Syst Biol 65: 726–736.

Holmes A. 2018. Analyzing Regulation of tRNAs, tRNA Fragments, and mRNAs in Whole Genomes. UC Santa Cruz https://escholarship.org/uc/item/4n09j1gw (Accessed August 19, 2019).

Hubisz MJ, Pollard KS, Siepel A. 2011. PHAST and RPHAST: phylogenetic analysis with space/time models. Brief Bioinform 12: 41–51.

Hummel G, Warren J, Drouard L. 2019. The multi-faceted regulation of nuclear tRNA gene transcription. IUBMB Life 71: 1099–1108. http://dx.doi.org/10.1002/iub.2097.

Johansson L, Gafvelin G, Arnér ESJ. 2005. Selenocysteine in proteins—properties and biotechnological use. Biochimica et Biophysica Acta (BBA) - General Subjects 1726: 1–13.

Jungreis I, Lin MF, Spokony R, Chan CS, Negre N, Victorsen A, White KP, Kellis M. 2011. Evidence of abundant stop codon readthrough in Drosophila and other metazoa. Genome Res 21: 2096–2113.

Kirchner S, Ignatova Z. 2015. Emerging roles of tRNA in adaptive translation, signalling dynamics and disease. Nat Rev Genet 16: 98–112.

Koski RA, Clarkson SG, Kurjan J, Hall BD, Smith M. 1980. Mutations of the yeast SUP4 tRNATyr locus: transcription of the mutant genes in vitro. Cell 22: 415–425.

Krinner S, Heitzer AP, Diermeier SD, Obermeier I, Längst G, Wagner R. 2014. CpG domains downstream of TSSs promote high levels of gene expression. Nucleic Acids Res 42: 3551–3564.

Kumar S, Stecher G, Suleski M, Hedges SB. 2017. TimeTree: A Resource for Timelines, Timetrees, and Divergence Times. Mol Biol Evol 34: 1812–1819.

Kutter C, Brown GD, Gonçalves A, Wilson MD, Watt S, Brazma A, White RJ, Odom DT. 2011. Pol III binding in six mammals shows conservation among amino acid isotypes despite divergence among tRNA genes. Nat Genet 43: 948–955.

Li WH, Ellsworth DL, Krushkal J, Chang BH, Hewett-Emmett D. 1996. Rates of nucleotide substitution in primates and rodents and the generation-time effect hypothesis. Mol Phylogenet Evol 5: 182–187.

Lorenz R, Bernhart SH, Höner Zu Siederdissen C, Tafer H, Flamm C, Stadler PF, Hofacker IL. 2011. ViennaRNA Package 2.0. Algorithms Mol Biol 6: 26.

Loughran G, Jungreis I, Tzani I, Power M, Dmitriev RI, Ivanov IP, Kellis M, Atkins JF. 2018. Stop codon readthrough generates a C-terminally extended variant of the human vitamin D receptor with reduced calcitriol response. J Biol Chem 293: 4434–4444.

Maraia RJ, Kenan DJ, Keene JD. 1994. Eukaryotic transcription termination factor La mediates transcript release and facilitates reinitiation by RNA polymerase III. Mol Cell Biol 14: 2147–2158.

Meunier J, Lemoine F, Soumillon M, Liechti A, Weier M, Guschanski K, Hu H, Khaitovich P, Kaessmann H. 2013. Birth and expression evolution of mammalian microRNA genes. Genome Res 23: 34–45.

Mleczko AM, Celichowski P, Bąkowska-Żywicka K. 2014. Ex-translational function of tRNAs and their fragments in cancer. Acta Biochim Pol 61: 211–216.

Necsulea A, Kaessmann H. 2014. Evolutionary dynamics of coding and non-coding transcriptomes. Nat Rev Genet 15: 734–748.

Necsulea A, Soumillon M, Warnefors M, Liechti A, Daish T, Zeller U, Baker JC, Grützner F, Kaessmann H. 2014. The evolution of lncRNA repertoires and expression patterns in tetrapods. Nature 505: 635–640.

Nguyen N, Hickey G, Zerbino DR, Raney B, Earl D, Armstrong J, Kent WJ, Haussler D, Paten B. 2015. Building a pan-genome reference for a population. J Comput Biol 22: 387–401.

Orioli A, Pascali C, Quartararo J, Diebel KW, Praz V, Romascano D, Percudani R, van Dyk LF, Hernandez N, Teichmann M, et al. 2011. Widespread occurrence of non-canonical transcription termination by human RNA polymerase III. Nucleic Acids Res 39: 5499–5512.

Palazzo AF, Lee ES. 2015. Non-coding RNA: what is functional and what is junk? Front Genet 6: 2.

Pan T. 2018. Modifications and functional genomics of human transfer RNA. Cell Res 28: 395–404.

Paten B, Diekhans M, Earl D, John JS, Ma J, Suh B, Haussler D. 2011a. Cactus graphs for genome comparisons. J Comput Biol 18: 469–481.

Paten B, Earl D, Nguyen N, Diekhans M, Zerbino D, Haussler D. 2011b. Cactus: Algorithms for genome multiple sequence alignment. Genome Res 21: 1512–1528.

Pedregosa F, Varoquaux G, Gramfort A, Michel V, Thirion B, Grisel O, Blondel M, Prettenhofer P, Weiss R, Dubourg V, et al. 2011. Scikit-learn: Machine Learning in Python. J Mach Learn Res 12: 2825–2830.

Pollard KS, Hubisz MJ, Rosenbloom KR, Siepel A. 2010. Detection of nonneutral substitution rates on mammalian phylogenies. Genome Res 20: 110–121.

Raab JR, Chiu J, Zhu J, Katzman S, Kurukuti S, Wade PA, Haussler D, Kamakaka RT. 2012. Human tRNA genes function as chromatin insulators. EMBO J 31: 330–350.

Roadmap Epigenomics Consortium, Kundaje A, Meuleman W, Ernst J, Bilenky M, Yen A, Heravi-Moussavi A, Kheradpour P, Zhang Z, Wang J, et al. 2015. Integrative analysis of 111 reference human epigenomes. Nature 518: 317–330.

Rogers HH, Bergman CM, Griffiths-Jones S. 2010. The evolution of tRNA genes in Drosophila. Genome Biol Evol 2: 467–477.

Roy B, Leszyk JD, Mangus DA, Jacobson A. 2015. Nonsense suppression by near-cognate tRNAs employs alternative base pairing at codon positions 1 and 3. Proc Natl Acad Sci U S A 112: 3038–3043.

Schaffer AE, Eggens VRC, Caglayan AO, Reuter MS, Scott E, Coufal NG, Silhavy JL, Xue Y, Kayserili H, Yasuno K, et al. 2014. CLP1 founder mutation links tRNA splicing and maturation to cerebellar development and neurodegeneration. Cell 157: 651–663.

Schmitt BM, Rudolph KLM, Karagianni P, Fonseca NA, White RJ, Talianidis I, Odom DT, Marioni JC, Kutter C. 2014. High-resolution mapping of transcriptional dynamics across tissue development reveals a stable mRNA-tRNA interface. Genome Res 24: 1797–1807.

Sun C, Fu Z, Wang S, Li J, Li Y, Zhang Y, Yang F, Chu J, Wu H, Huang X, et al. 2018. Roles of tRNA-derived fragments in human cancers. Cancer Lett 414: 16–25.

Tang DTP, Glazov EA, McWilliam SM, Barris WC, Dalrymple BP. 2009. Analysis of the complement and molecular evolution of tRNA genes in cow. BMC Genomics 10: 188.

Thornlow B, Hough J, Roger J, Gong H, Lowe T, Corbett-Detig R. 2018. Transfer RNA genes experience exceptionally elevated mutation rates. Proceedings of the National Academy of Sciences 115: 8996–9001.

Valle RP, Morch MD, Haenni AL. 1987. Novel amber suppressor tRNAs of mammalian origin. EMBO J 6: 3049–3055.

Yacoubi BE, El Yacoubi B, Bailly M, de Crécy-Lagard V. 2012. Biosynthesis and Function of Posttranscriptional Modifications of Transfer RNAs. Annual Review of Genetics 46: 69–95. http://dx.doi.org/10.1146/annurev-genet-110711-155641.

Yeganeh M, Praz V, Cousin P, Hernandez N. 2017. Transcriptional interference by RNA polymerase III affects expression of the Polr3e gene. Genes & Development 31: 413–421. http://dx.doi.org/10.1101/gad.293324.116.

Yoo H, Son D, Jang Y-J, Hong K. 2016. Indispensable role for mouse ELP3 in embryonic stem cell maintenance and early development. Biochem Biophys Res Commun 478: 631–636.

Zheng G, Qin Y, Clark WC, Dai Q, Yi C, He C, Lambowitz AM, Pan T. 2015. Efficient and quantitative high-throughput tRNA sequencing. Nat Methods 12: 835–837.

