## Supplemental_Material for "Predicting transfer RNA gene activity from sequence and genome context"

**Caveats to tRNA gene classification methods.** Ideally, we would observe that all species contained at least one tRNA gene per expected anticodon (excluding anticodons for which no active tRNA gene is observed in human or mouse). We found three exceptions to this rule. *C. hircus* (goat) is the only species without a predicted active tRNA-Ser-TGA gene. *C. hircus* is also one of only three species that has any tRNA-Ser-GGA genes, and it has 27 such genes, one of which was predicted active. However, these most likely represent assembly errors, as 21 of these share a 10-base 3' flanking motif (AGAAGGAAAT), and Ser-GGA is not a standard eukaryotic tRNA (Grosjean et al. 2010). We also predict that the lone tRNA-Leu-TAA gene in *Erinaceus europaeus* (hedgehog) is inactive. The probability score for this prediction is 0.51, nearly the minimum possible score, indicating that it may be active. Additionally, the *Chinchilla lanigera* (chinchilla) genome in our Cactus graph (ChiLan1.0) does not contain any high-confidence selenocysteine (SeC) tRNA genes. This is likely due to an incomplete genome assembly, as selenocysteine tRNAs are required in mammals (Johansson et al. 2005).

We also identified several anomalous lineage-specific expansions that may represent SINEs or assembly errors. Notably, *E. europaeus* has 123 tRNA-Lys-CTT genes. All have high tRNAscan-SE general bit scores (Supplemental Fig. S10) and are unlikely to be pseudogenes. We classified 98 of these as active. tRNA-derived SINEs can proliferate rapidly via retrotransposition, and a rapid expansion may have occurred in this species, targeting areas within or near regions of high transcription. While few of these tRNA genes have completely identical sequences, 43 have identical 3' flanking regions, extending 10 bases after the tRNA gene (CCATTTGTTG). An additional 24 genes share a different 3' flanking motif (CCAGATGTTG). The immediate flanking regions of actively transcribed tRNA genes are expected to be highly divergent due to transcription-associated mutagenesis (Thornlow et al. 2018). One possible explanation for this discrepancy is that these motifs are part of a fused SINE element responsible for the proliferation of the tRNA-derived SINE element. A similar expansion has occurred in *Microtus ochrogaster* (prairie vole), whose genome contains 131 tRNA-Lys-CTT genes, 66 of which we classified as active. 25 of these genes are exactly identical in tRNA gene sequence, suggesting a similarly recent expansion. These difficulties illuminate a benefit of our classifier in increased detection of SINE elements, as each of these genes passes through the high confidence filter in tRNAscan-SE, but may not represent truly actively transcribed tRNA loci.

**Identification of a possibly active chimpanzee nonsense suppressor tRNA gene.** To our knowledge, no actively transcribed nonsense suppressor tRNA gene has been demonstrated in a primate genome. However, we have predicted a tRNA-Sup-CTA in *Pan troglodytes* (chimpanzee) as active, although experimentation is necessary to support that this tRNA gene is indeed a suppressor and is also actively transcribed. While nonsense suppression via readthrough by near-cognate tRNAs has been demonstrated several times in metazoa (Jungreis et al. 2011; Yacoubi et al. 2012; Loughran et al. 2018; Roy et al. 2015) and suppressor tRNAs have been found in other species (Beier and Grimm 2001; Valle et al. 1987), evidence for an actively transcribed suppressor tRNA in a primate species is of great interest for future experimentation. This tRNA gene is a conserved ortholog of the human tRNA-Gln-CTG-6-1 gene, and exhibits several chimpanzee-specific nucleotide substitutions (Supplemental Fig. S11) that are not expected to significantly impair the function of its transcript. However, this finding may merely reflect an assembly error. In some chimpanzee assemblies, this is a tRNA-Gln-CTG gene while in others it is a tRNA-Sup-CTA gene. Therefore, this gene may in fact be an actively transcribed chimpanzee tRNA-Gln-CTG gene.

### SUPPLEMENTAL METHODS:

**Generating feature data.** To generate PhyloP data, we used the HAL (Hickey et al. 2013) and PHAST (Hubisz et al. 2011) toolkits. We extracted four-fold degenerate (4d) sites from each genome of interest with the hal4dExtract command, reduced each set of 4d sites to 100,000 sites, and trained a PhyloP model with this reduced set using the phyloFit command. We then extracted each genome of interest from the Cactus graph using the hal2fasta command, and applied tRNAscan-SE 2.0 (Chan et al. 2019) to each using the -o, -f, -s, -m, -b, -a, --detail, -H, and -y flags. We applied EukHighConfidenceFilter using the -r, -i, -s, -p, and -o flags. For classification purposes, we used all tRNAs in the data output by the EukHighConfidenceFilter, which includes high-confidence tRNA genes, as well as tRNA genes with high tRNAscan-SE bit scores and isotype mismatches. From there, we created custom Python scripts (available at https://github.com/bpt26/tRNA_classifier) to generate .bed files and .fasta files including tRNA loci and their flanking regions, up to 350 nucleotides upstream and downstream of each gene. We applied the hal2maf and PhyloP commands to generate PhyloP data for these loci, as well as the phastCons command to identify tRNA genes with conserved flanking regions. For conserved elements no larger than 100 base pairs with phastCons scores of at least 200, we used the boundaries of the conserved element instead of the tRNA boundaries, as conserved flanking regions might reduce the accuracy of classification. We applied this process to obtain all feature data for all species, including human. To determine performance on the human data, we used ten-fold cross-validation. For all other species, we trained a random forest classifier using ten-fold cross-validation in scikit-learn on the human data, and then applied it to feature data for each species of interest. Our final model in scikit-learn uses 250 estimators, a minimum of 2 samples required to split an internal node and a maximum depth of 4 nodes (Pedregosa et al. 2011).

To estimate the minimum free energy of canonical secondary structure, we used folding constraints output by tRNAscan-SE (Chan et al. 2019) as inputs to RNAfold (Lorenz et al. 2011), which estimates minimum free energy. Initially, we examined both RNAfold and tRNAscan-SE output. RNAfold output is significantly correlated with tRNAscan-SE secondary structure scores (Spearman rank correlation, $p < 2e-4$), indicating interchangeability. To determine which to include in the model, we created a separate model that included both scores, and found that RNAfold output had greater feature importance score (0.069 for RNAfold output compared to 0.013 for tRNAscan-SE secondary structure score). We also tested a model that includes tRNAscan-SE secondary structure, HMM, and isotype-specific bit scores. Upon their inclusion, the feature importance score of the tRNAscan-SE general bit score decreases slightly from 0.055 to 0.048, but it remains more important to the model than any of the other tRNAscan-SE scores (secondary structure, HMM and isotype-specific bit scores had feature importance values of .012, .035 and .041, respectively), supporting its usage in the model.

**Fitting to a Markov Model.** For each species, we converted our ortholog set to an alignment, where for each species, ‘2’ indicated a predicted active tRNA, ‘1’ indicated a predicted inactive tRNA, and ‘0’ indicated no detected ortholog. We used RevBayes (Höhna et al. 2016) to fit this data, in conjunction with a phylogeny from TimeTree (Kumar et al. 2017), to a Markov model and allowed only the Q matrix to be changed over 10,000 generations. We then used the getTransitionProbabilities() command to estimate transition probabilities to and from each state over branch lengths of 10 million years.

The guide tree for the Cactus graph was:

((((((((Homo_sapiens:0.00655,Pan_troglodytes:0.00684)Anc32:0.00422,Gorilla_gorilla_gorilla:0.008964)Anc29:0.009693,Pongo_abelii:0.01894)Anc25:0.015511,Macaca_mulatta:0.043601)Anc20:0.08444,Aotus_nancymaae:0.08)Anc15:0.08,Microcebus_murinus:0.10612)Anc11:0.083494,((((Jaculus_jaculus:0.1,(Microtus_ochrogaster:0.14,(Mus_musculus:0.084509,Rattus_norvegicus:0.091589)Anc30:0.047773)Anc26:0.06015)Anc21:0.122992,(Heterocephalus_glaber:0.1,(Cavia_porcellus:0.065629,(Chinchilla_lanigera:0.06,Octodon_degus:0.1)Anc31:0.06)Anc27:0.05)Anc22:0.06015)Anc16:0.05,Marmota_marmota:0.1)Anc12:0.05,Oryctolagus_cuniculus:0.21569)Anc08:0.04)Anc04:0.040593,(((Sus_scrofa:0.12,(Orcinus_orca:0.069688,(Bos_taurus:0.04,Capra_hircus:0.04)Anc23:0.09)Anc17:0.045488)Anc13:0.02,((Equus_caballus:0.109397,(Felis_catus:0.098612,(Canis_lupus_familiaris:0.052458,Mustela_putorius_furo:0.08)Anc28:0.02)Anc24:0.049845)Anc18:0.02,(Pteropus_alecto:0.1,Eptesicus_fuscus:0.08)Anc19:0.033706)Anc14:0.03)Anc09:0.025,Erinaceus_europaeus:0.278178)Anc05:0.021227)Anc02:0.023664,((Loxodonta_africana:0.098842,Chrysochloris_asiatica:0.04)Anc06:0.05,Dasypus_novemcinctus:0.169809)Anc03:0.02)backbone_root;

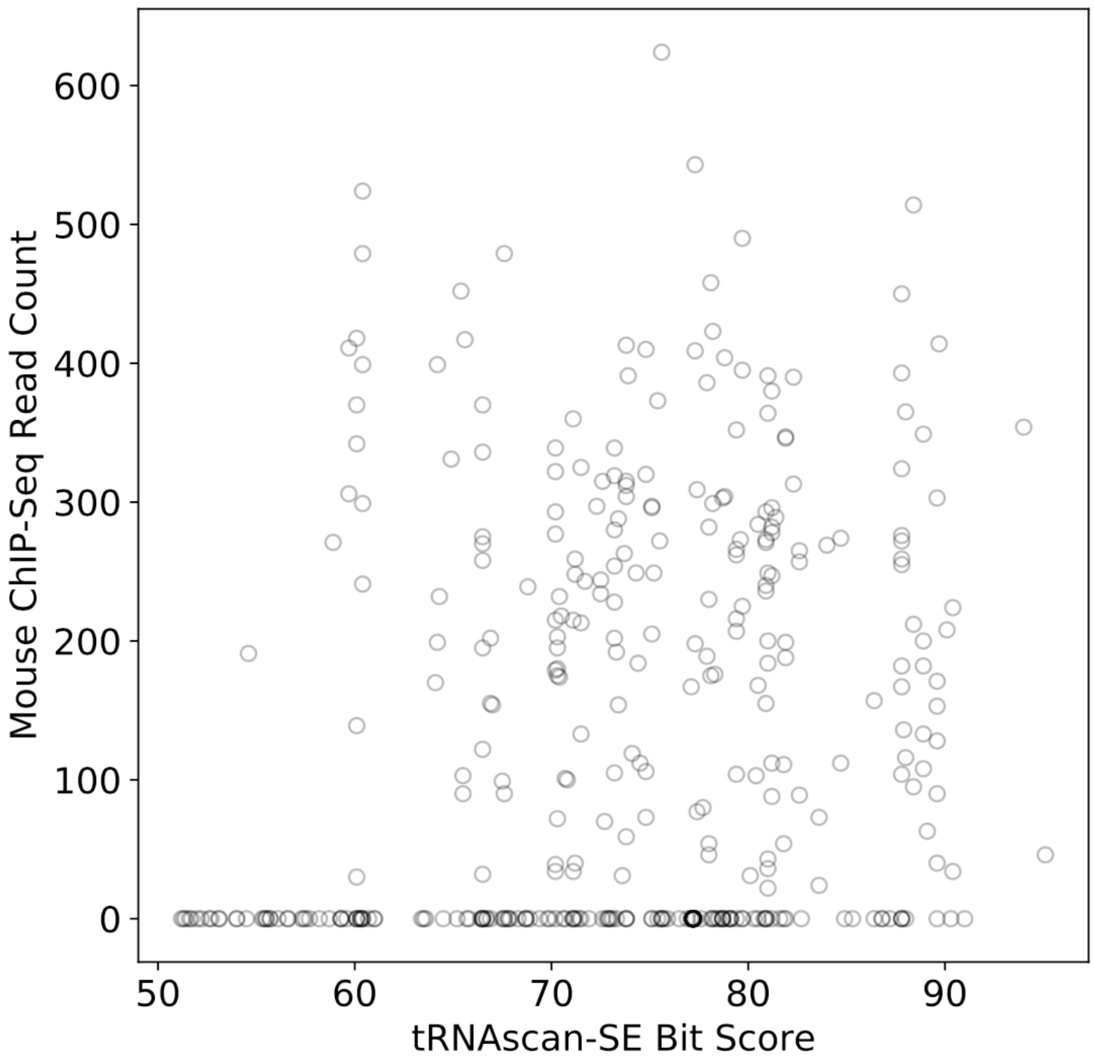


**Supplemental Figure S1: tRNAscan-SE general bit score alone is not sufficient for inferring transcriptional activity.** tRNAscan-SE general bit scores are compared to the highest RNA Polymerase III ChIP-Seq read count across mouse liver, muscle and testes for each mouse tRNA gene (Kutter et al. 2011). Many relatively high-scoring tRNA genes (greater than ~70 bits) have no evidence for activity.

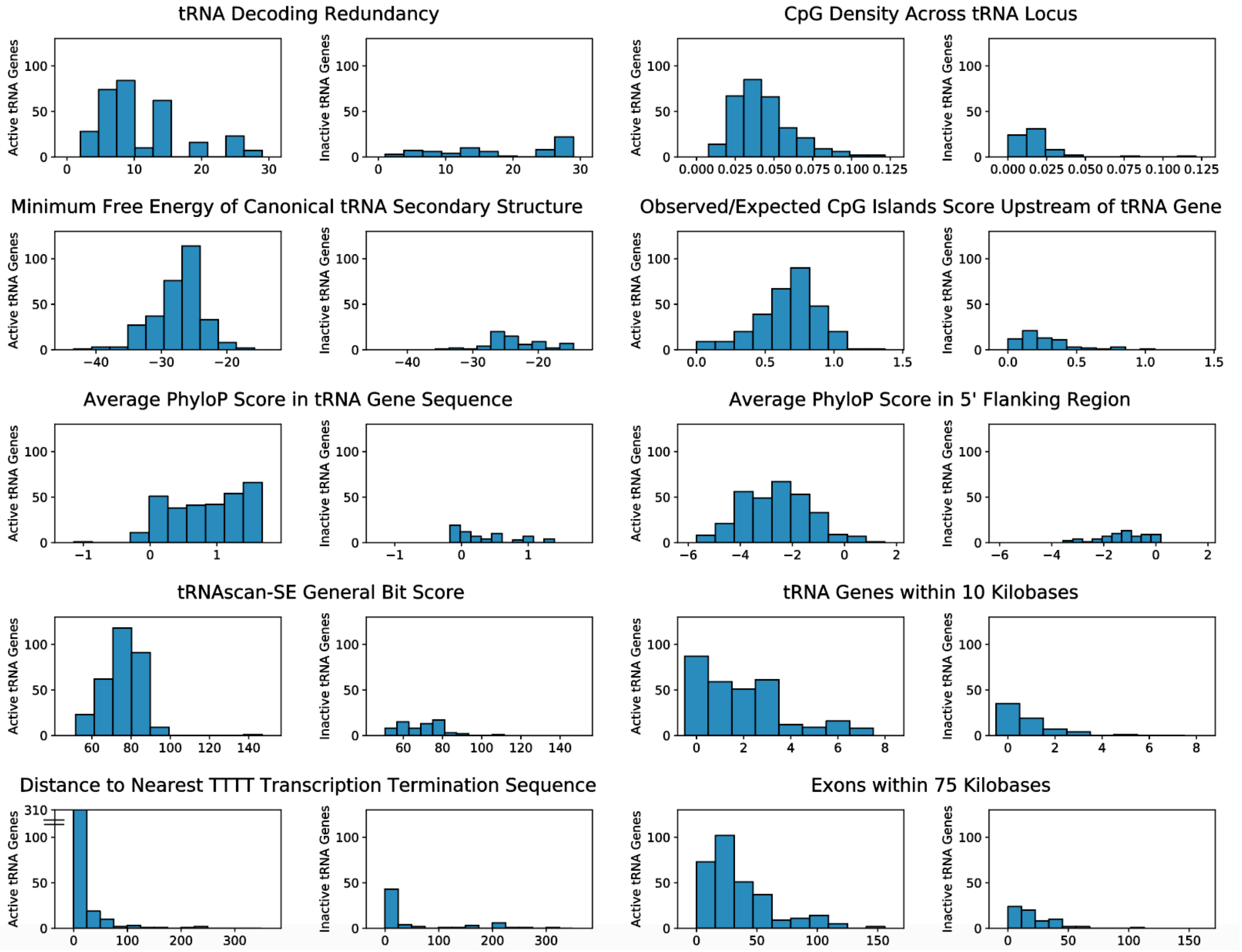


**Supplemental Figure S2: Distributions of all features for active and inactive tRNA genes in the training set.** For all intrinsic (left) and extrinsic (right) features in our model, the distribution across all active and inactive tRNA genes is shown. The mean and standard deviation for each feature can be found in Supplemental Table S2.

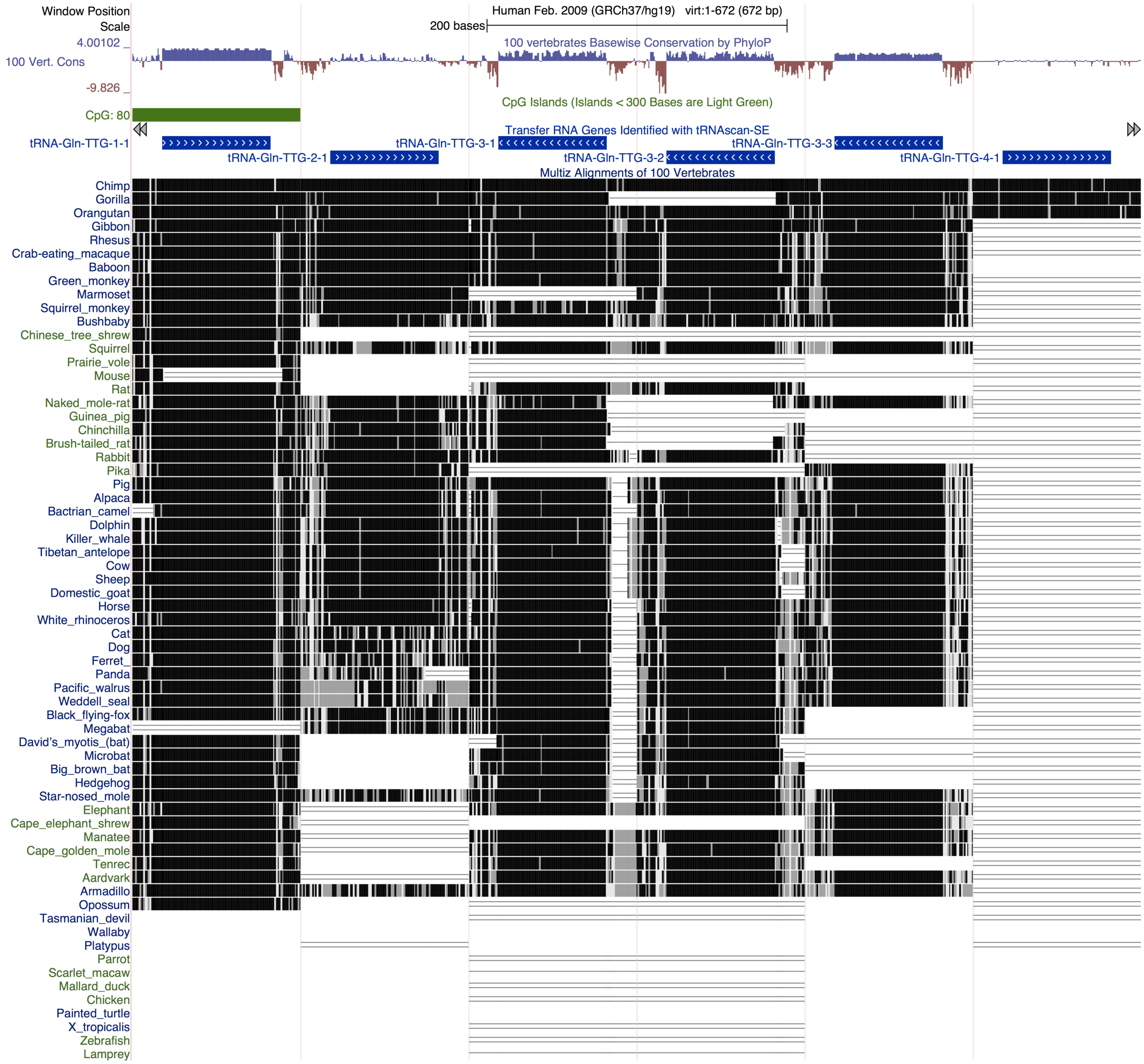


**Supplemental Figure S3: Visualization of tRNA-Gln-TTG genes in the UCSC Genome Browser.** From Table 1, all six tRNA-Gln-TTG genes, flanked by 20 nucleotides on either side, in the human genome (GRCh37) are shown with the PhyloP track (Pollard et al. 2010; labeled 100 Vert. Cons), CpG Islands track (Gardiner-Garden and Frommer 1987; labeled CpG) and the Multiz alignment tracks (Blanchette et al. 2004; bottom). Blue regions in the PhyloP track indicate sequence conservation and brown regions indicate accelerated evolution. Green blocks in the CpG islands track indicate regions enriched for CpG dinucleotides. Black regions in the Multiz alignment indicate alignments to genomic regions in the indicated species. Differences in PhyloP scores are due to the use of alignments across 100 vertebrates in the track shown, while our study used 29-way eutherian alignments.

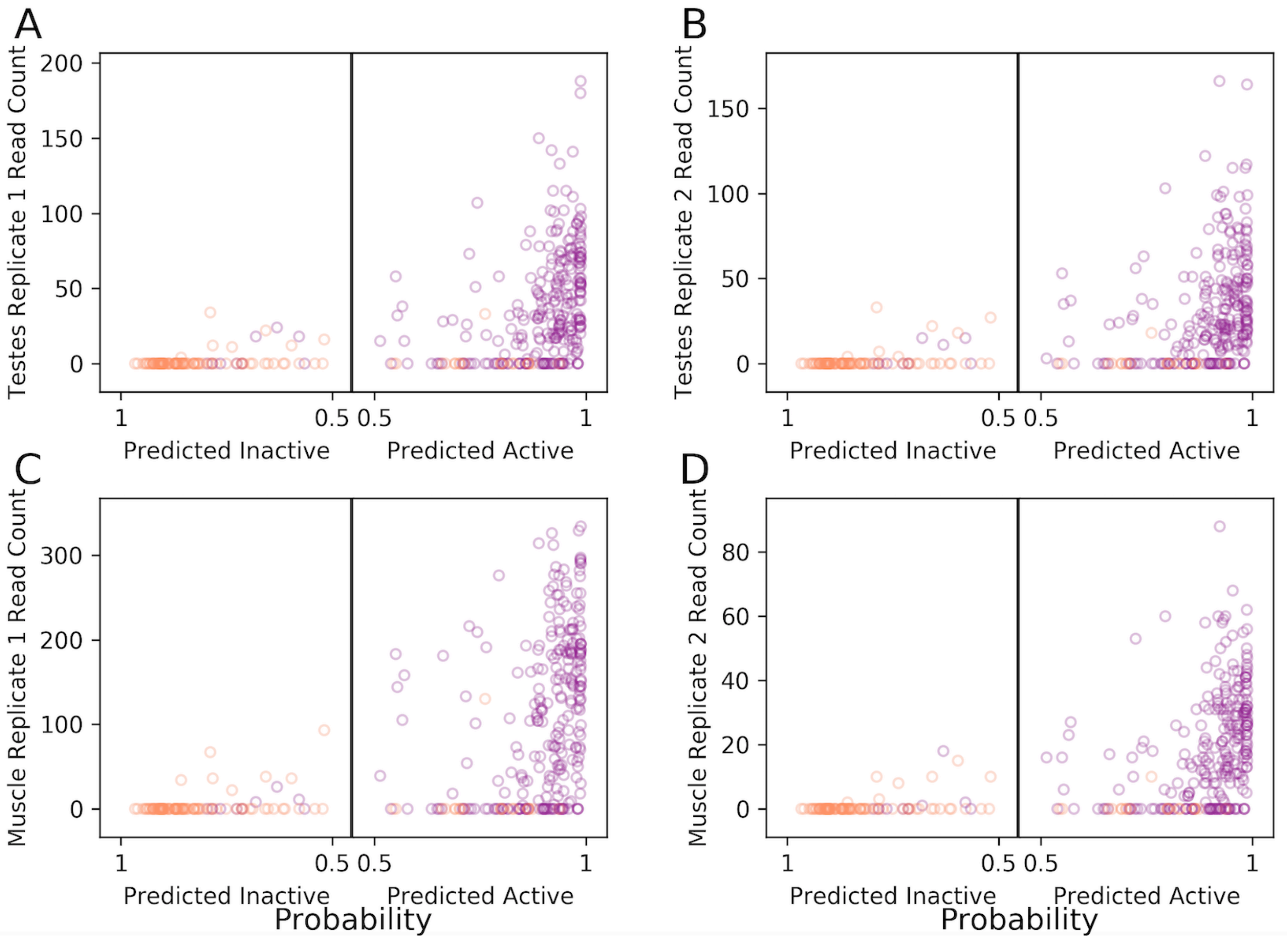


**Supplemental Figure S4: Comparison of tRNA gene classifications to ChIP-Seq data is consistent across tissues.** Mouse ChIP-Seq data compared to tRNA gene classifications and ChIP data for both testes replicates and both muscle replicates. Probability scores output by the classifier are shown on the horizontal axis where tRNA genes furthest left are predicted inactive with greater probability and tRNA genes furthest right are predicted active with greater probability. The vertical axis indicates read counts in (A, B) testes and (C, D) muscle from Kutter et al. 2011. Purple indicates measured activity in at least one tissue based on chromatin modification data from Bogu et al. 2015 while orange indicates no measured activity in any tissues.

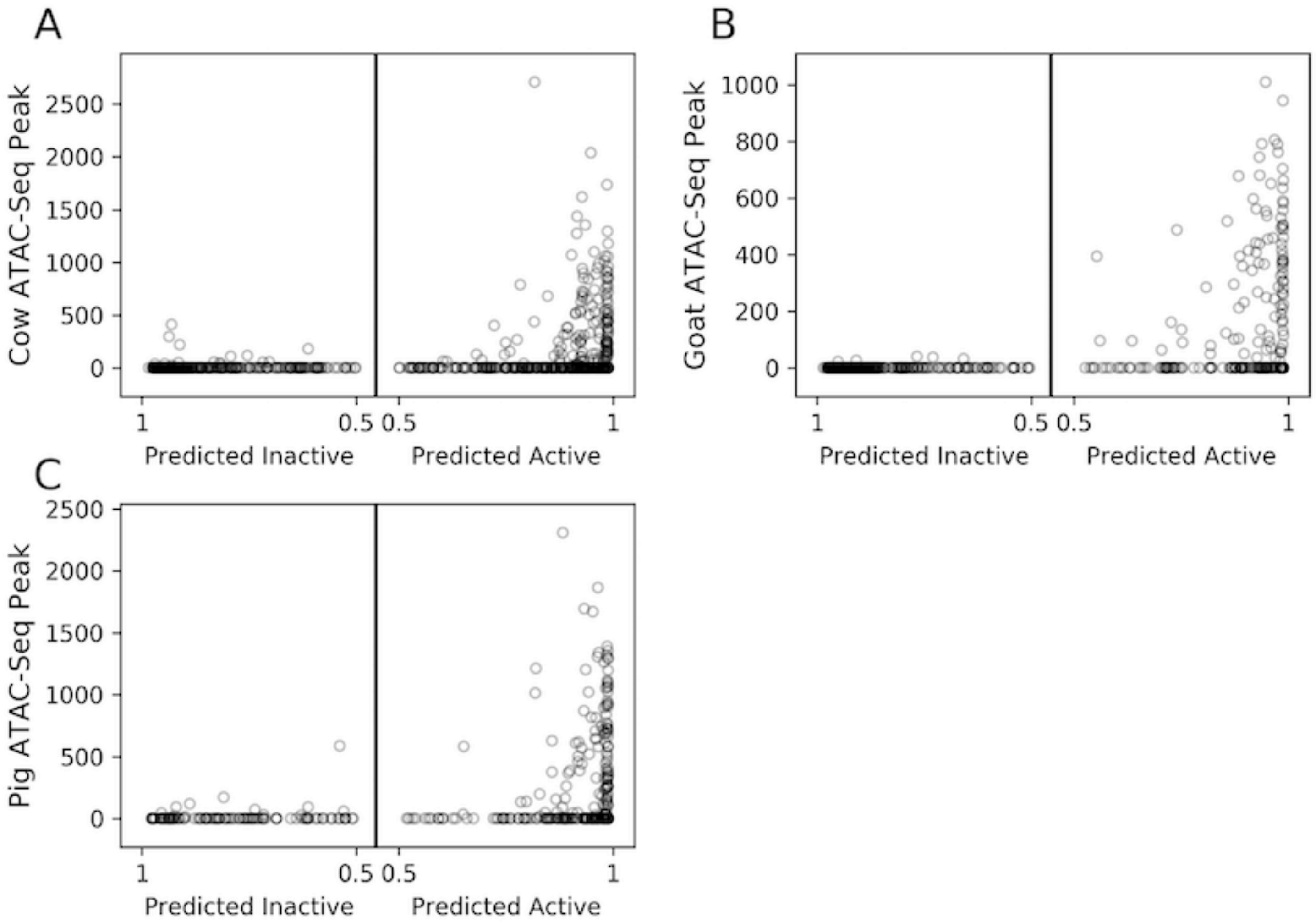


**Supplemental Figure S5: Comparison of classifications to ATAC-Seq data suggests similar accuracy across clades.** The mean ATAC-Seq peaks across liver, CD4 and CD8 cells in cow (A), goat (B) and pig (C) within 250 base pairs of each tRNA gene are shown against the classification probabilities for each gene as in Fig. 3.

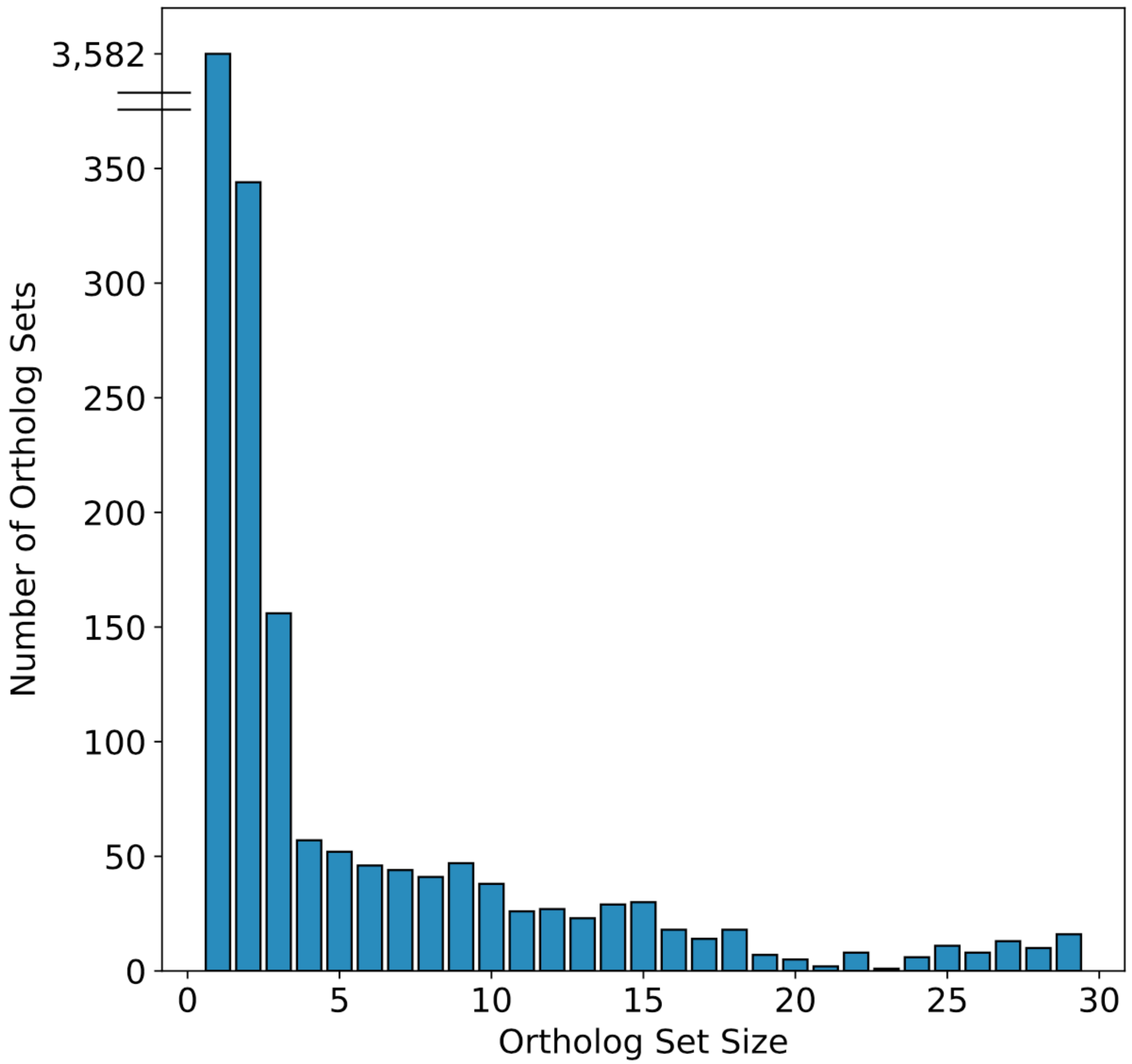


**Supplemental Figure S6: tRNA genes are generally either fairly deeply conserved or recently evolved.** Size distribution of all ortholog sets, with genes present in at least 2 species. 3,582 of the 11,752 tRNA genes in our alignment are species-specific. The remaining 8,170 tRNA genes are condensed into 1,097 ortholog sets, ranging in size from 2 to 29 species.

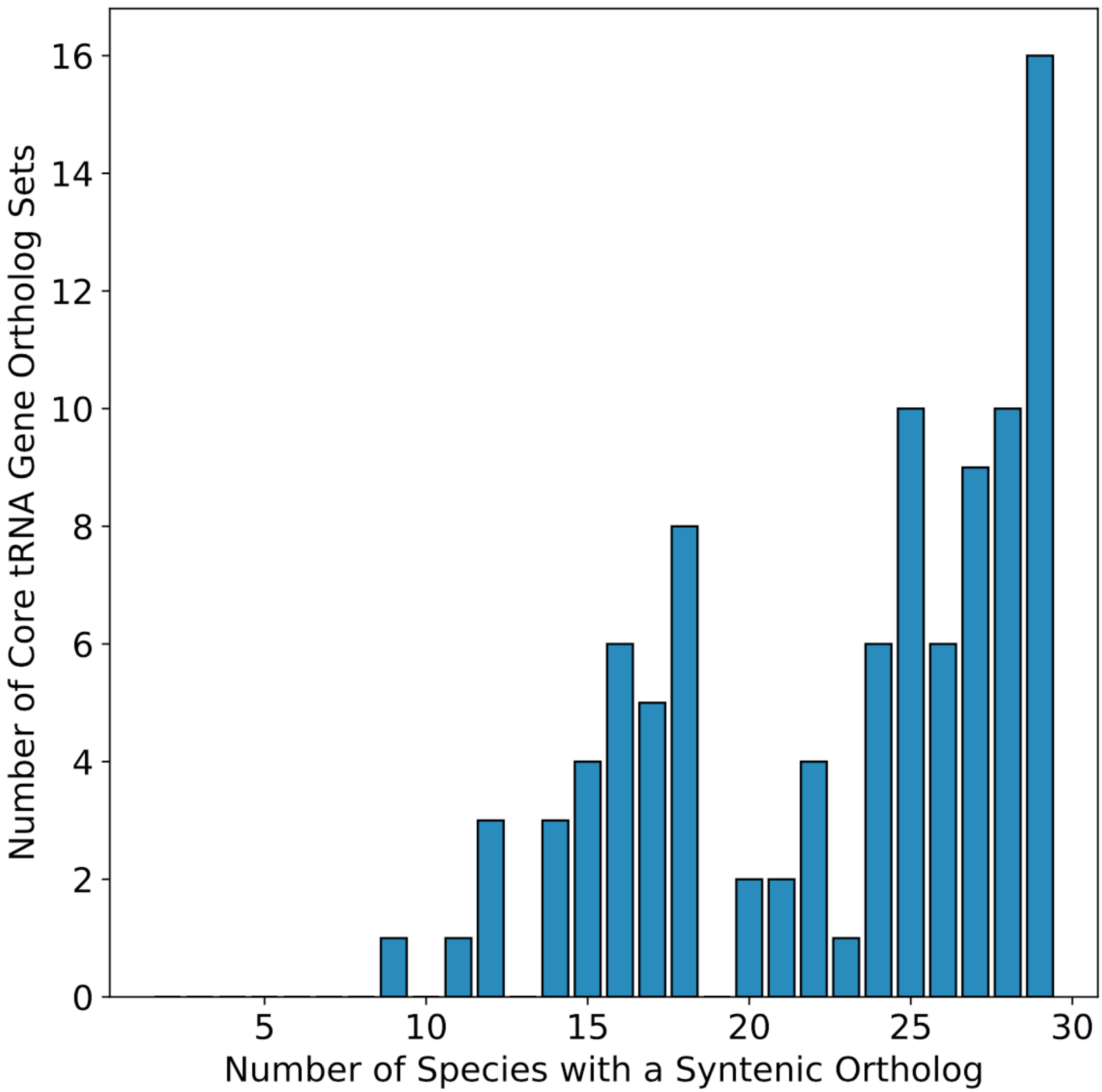


**Supplemental Figure S7: Core tRNA genes are very deeply conserved.** Histogram of sizes of the 97 “core” tRNA gene ortholog sets present in at least all 7 primate species.

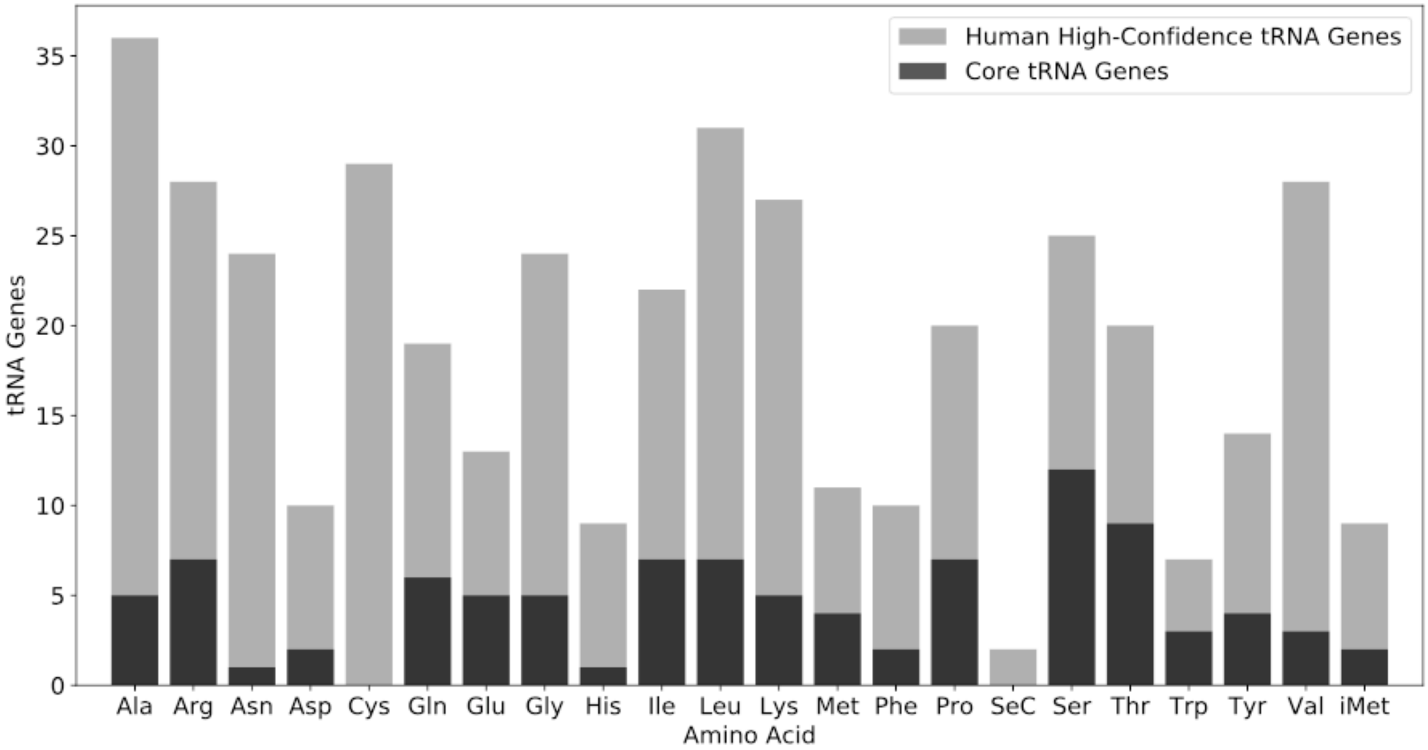


**Supplemental Figure S8: Cysteine is the only standard amino acid without a "core" tRNA.** All human active and inactive high-confidence tRNA genes are shown in gray for each amino acid, and those found in the core set of 97 tRNA genes are shown in black. Cysteine is one of the highest copy-number tRNA gene families, but is the only standard amino acid with zero "core" tRNA genes.

**A**

| Initial | Final: Missing | Inactive | Active |
|---|---|---|---|
| Missing | .871 | .111 | .017 |
| Inactive | .092 | .859 | .049 |
| Active | .012 | .014 | .974 |

**B**

| Initial | Final: Missing | Inactive | Active |
|---|---|---|---|
| Missing | .403 | .341 | .256 |
| Inactive | .116 | .845 | .040 |
| Active | .144 | .419 | .437 |

**C**

| Initial | Final: Missing | Inactive | Active |
|---|---|---|---|
| Missing | .723 | .229 | .049 |
| Inactive | .163 | .656 | .182 |
| Active | .106 | .365 | .528 |

**D**

| Initial | Final: Missing | Inactive | Active |
|---|---|---|---|
| Missing | .957 | .022 | .021 |
| Inactive | .165 | .821 | .015 |
| Active | .150 | .188 | .663 |

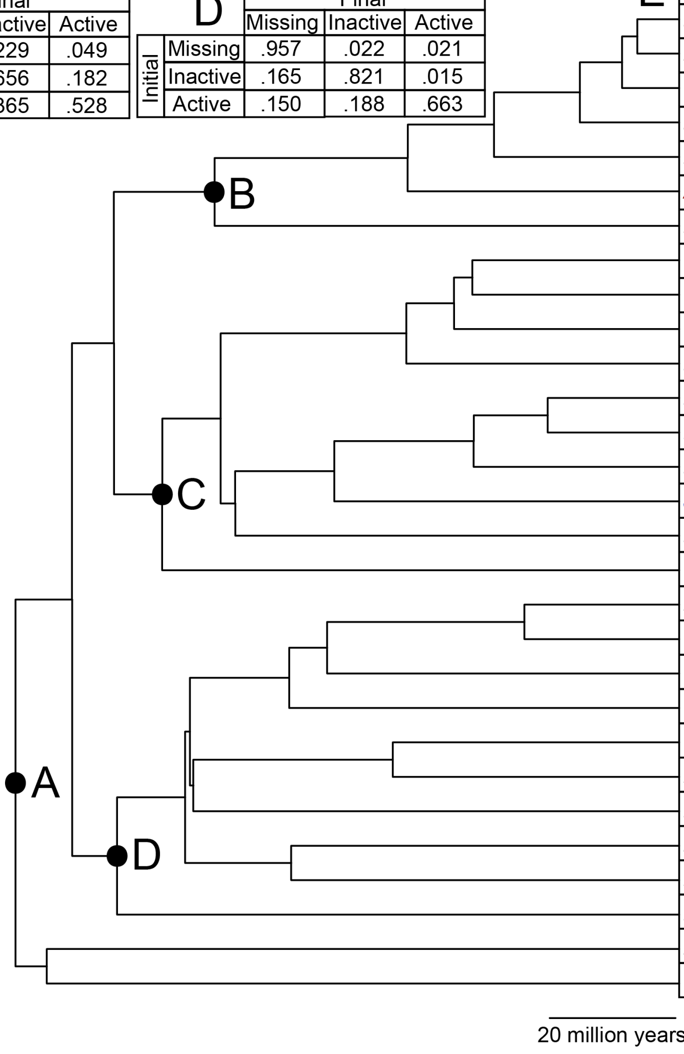


**E**

| Species | Total | High Conf. | % Active |
|---|---|---|---|
| *Homo sapiens* | 631 | 431 | 84.9 |
| *Pan troglodytes* | 590 | 399 | 85.5 |
| *Gorilla gorilla gorilla* | 566 | 375 | 87.7 |
| *Pongo abelii* | 618 | 411 | 87.8 |
| *Macaca mulatta* | 747 | 393 | 86.3 |
| *Aotus nancymaae* | 535 | 334 | 87.2 |
| *Microcebus murinus* | 467 | 328 | 87.8 |
| *Octodon degus* | 144,501 | 298 | 80.5 |
| *Chinchilla lanigera* | 51,076 | 301 | 83.1 |
| *Cavia porcellus* | 74,328 | 350 | 79.4 |
| *Heterocephalus glaber* | 28,957 | 294 | 81.0 |
| *Mus musculus* | 40,912 | 408 | 70.3 |
| *Rattus norvegicus* | 211,167 | 381 | 71.7 |
| *Microtus ochrogaster* | 29,471 | 470 | 73.0 |
| *Jaculus jaculus* | 119,136 | 318 | 77.4 |
| *Marmota marmota* | 188,473 | 411 | 61.3 |
| *Oryctolagus cuniculus* | 4,509 | 483 | 85.5 |
| *Bos taurus* | 263,431 | 810 | 54.0 |
| *Capra hircus* | 210,557 | 547 | 41.7 |
| *Orcinus orca* | 176,548 | 667 | 49.8 |
| *Sus scrofa* | 63,225 | 433 | 76.0 |
| *Canis lupus familiaris* | 150,565 | 382 | 76.4 |
| *Mustela putorius furo* | 182,506 | 380 | 77.1 |
| *Equus caballus* | 1,970 | 419 | 84.5 |
| *Pteropus alecto* | 4,802 | 345 | 64.3 |
| *Eptesicus fuscus* | 102,092 | 214 | 93.0 |
| *Erinaceus europaeus* | 7,180 | 447 | 76.7 |
| *Dasypus novemcinctus* | 227,726 | 539 | 72.4 |
| *Chrysochloris asiatica* | 26,044 | 372 | 74.0 |

**Supplemental Figure S9: Clades vary in their activity state transition rates.** (A-D): Transition matrices shown as in Fig. 4 for all descendants of labeled nodes. Phylogeny, species names and predicted active and inactive tRNA gene counts reproduced from Fig. 4. (E): For each species in our phylogeny (Hedges et al. 2006), the number of tRNA genes annotated by tRNAscan-SE 2.0 (Total; Chan et al. 2019), the number of tRNA genes present after application of the high-confidence filter (High Conf.) and the percent of such tRNA genes classified as active (% Active), are shown. Primate species are colored in red and rodent species are colored in blue. In contrast to Fig. 4B, the High Conf. column includes tRNA genes within segmental duplications.

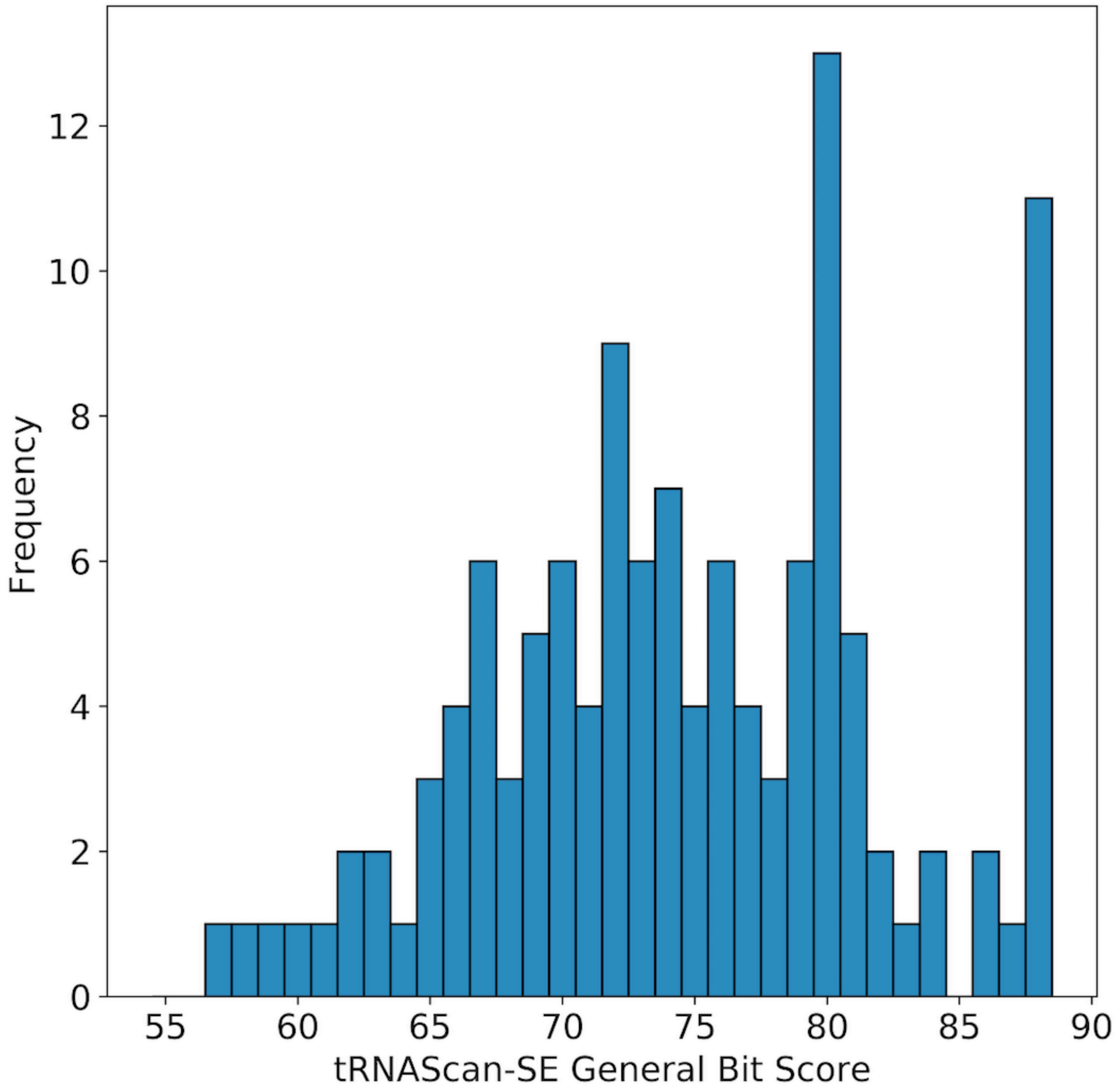


**Supplemental Figure S10: Most *Erinaceus europaeus* tRNA-Lys-CTT genes have high tRNAScan-SE general bit scores.** Histogram showing the number of tRNA-Lys-CTT genes in the *E. europaeus* genome by tRNAScan-SE general bit score. For reference, the average active human tRNA gene has a bit score of 76.2 and the average inactive human tRNA gene has a bit score of 69.3.

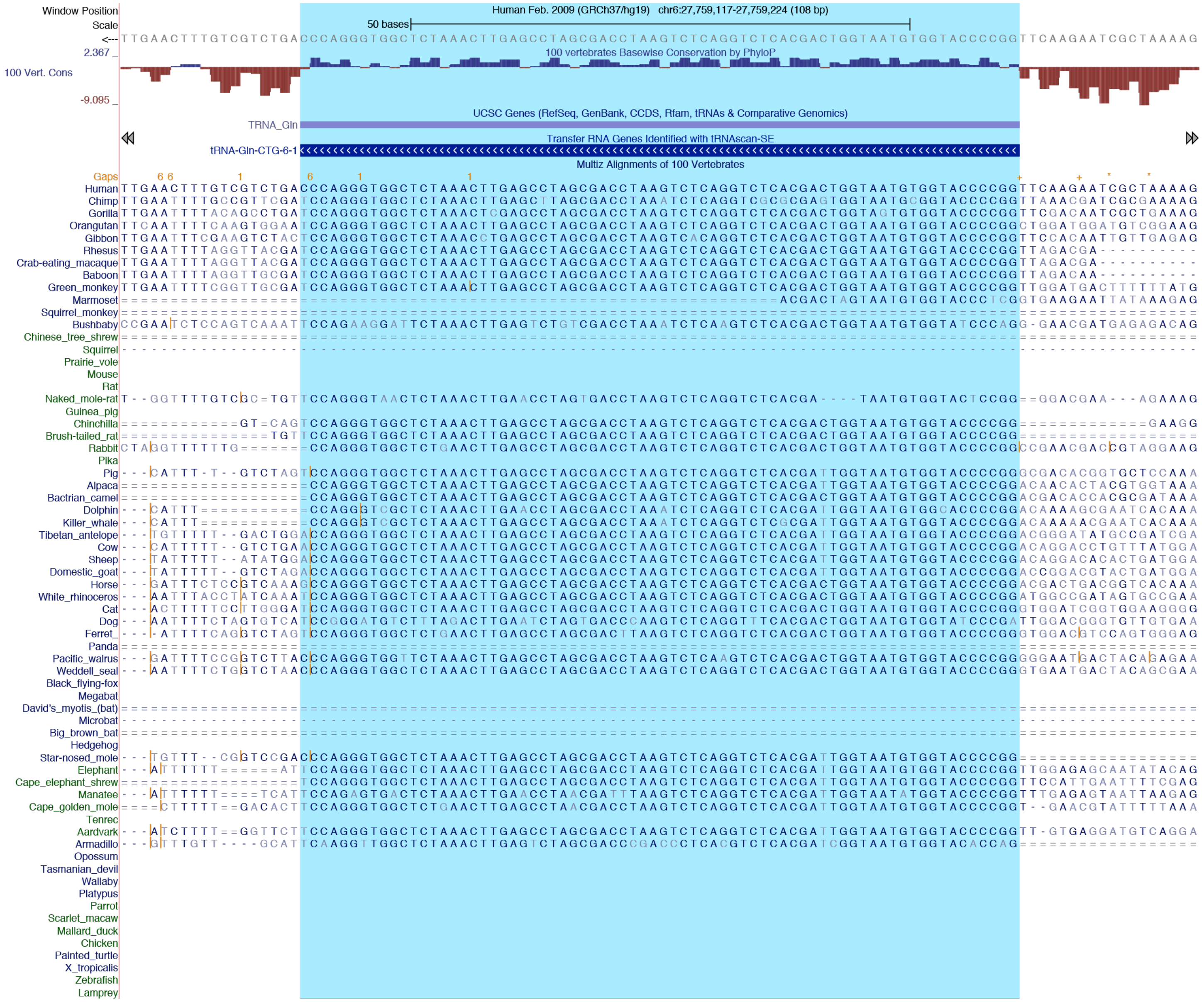

**Supplemental Figure S11: Chimpanzee ortholog of human tRNA-Gln-CTG-6-1 may be an actively transcribed suppressor tRNA gene.** Gene browser screenshot showing the PhyloP and Multiz Alignment tracks of the human tRNA-Gln-CTG-6-1 gene, for which the chimpanzee genome has a tRNA-Sup-CTA gene. The sequence of the tRNA gene is highlighted in cyan. The chimpanzee sequence (second line from top) exhibits several unique substitutions. Most notably, the CTG anticodon (here from right to left in the center of the highlighted region) has changed to CTA in chimp, accompanied by other changes to the body of the tRNA. The PhyloP track also shows strong divergence upstream and downstream of the tRNA gene sequence, consistent with high levels of expression.
